# A Supramolecular Self-assembly Approach to Site-Specific Antibody Conjugates *via* a Coiled-coil Peptides Platform

**DOI:** 10.1101/2025.07.21.665979

**Authors:** Alina Ringaci, Ting-Yu Shih, Mark W. Grinstaff

## Abstract

Antibody conjugates play a central role across multiple healthcare sectors with a prime example being antibody-drug conjugates (ADCs). Although widely used lysine and hinge cysteine conjugation methods yield products, the lack of site-specificity and spatial control along with the highly heterogeneous composition are significant limitations. We describe a facile supramolecular assembly method based on heterodimer coiled-coil formation for site-specific antibody conjugation. The method affords uniform loading of diverse payloads including anti-cancer agents, polymers, enzymes, fluorophores, etc. under mild aqueous conditions. Further, the facile convergent approach capitalizes on the independent strengths and flexibility of protein expression and peptide chemistry culminating in a final self-assembly step. Coiled-coil conjugation perseveres both antibody antigen binding sites for target engagement and heavy chains constant domains for Fc binding and recycling. An ADC loaded with monomethyl auristatin E targeting HER2+ tumors significantly reduces tumor volume in a human ovarian cancer xenograft model outperforming the antibody alone with validated performance against a best-in-class therapeutic. Supramolecular assembly-driven bioconjugation expands the bioorthogonal chemistry toolbox for antibody modification and opens new avenues for advanced antibody conjugates with multiple payloads.

## INTRODUCTION

Biorthogonal chemistry, the selective chemical transformation within a complex biological milieu (1,2), enables the modification of proteins to impart increased functionality and utility. The preparation of antibody-drug conjugates (ADCs) highlights a specific use case, and ADCs are transformative treatments against cancer. Classical and still widely used bioconjugation methods for ADCs include reacting protein primary amines with an entity containing an activated ester or converting primary amines to thiols, via Traut’s reagent, or using hinge cysteines for subsequent coupling to a maleimide functionalized moiety (3–5). Although they are useful reactions, their selectivity and specificity are poor with ill-defined, highly heterogeneous products being obtained from a single reaction (6,7). Newer techniques rely on unique reactive sites such as a C-terminal thioester or N-terminal amine and afford well-defined products (8,9). Additional significant methodology advancements include: 1) incorporation of a non-natural amino acid into the protein such as azide-modified phenylalanine (10–13) or glycan remodeling (14–16) for a subsequent specific click-chemistry reaction (e.g., azide fragment with an alkyne functionalized molecule); 2) disulfide rebridging (17,18); and, 3) enzymatic ligation reactions (e.g., AviTag, SorTag) (19–22) and split proteins tags (e.g. SpyTag, Intein Tag) (23–26). Each of these methods possess a number of advantages for the preparation of ADCs, but none have advanced to preparation of clinically approved products (27,28).

A supramolecular assembly synthesis strategy offers benefits such as ease of use, selectivity, bio-orthogonality, and complementarity to other methods, but is lacking from the current synthesis toolbox for antibody conjugates. Supramolecular assembly, the ability of smaller entities to spontaneously organize into well-defined larger structures through non-covalent interactions, is widely used in materials science, chemistry, and biological chemistry (29–33). Alpha-helical coil peptides are protein structural motifs capable of orthogonal and selective self-assembly into coiled-coil structures (**Figure 1A**). Although they rely on non-covalent interactions, the presence of multiple contact points within the assembly significantly contributes to stability with dissociation constants in the sub-nanomolar range (34). Due to modular design and easily controlled molecular features, by varying the peptide sequences, coiled-coils are used in many biomedical applications (35–38) including hydrogel fabrication (39,40), nanoparticle modification (41,42), fusion protein preparation (43,44), and cell therapy engineering (45,46).

**Figure 1.**
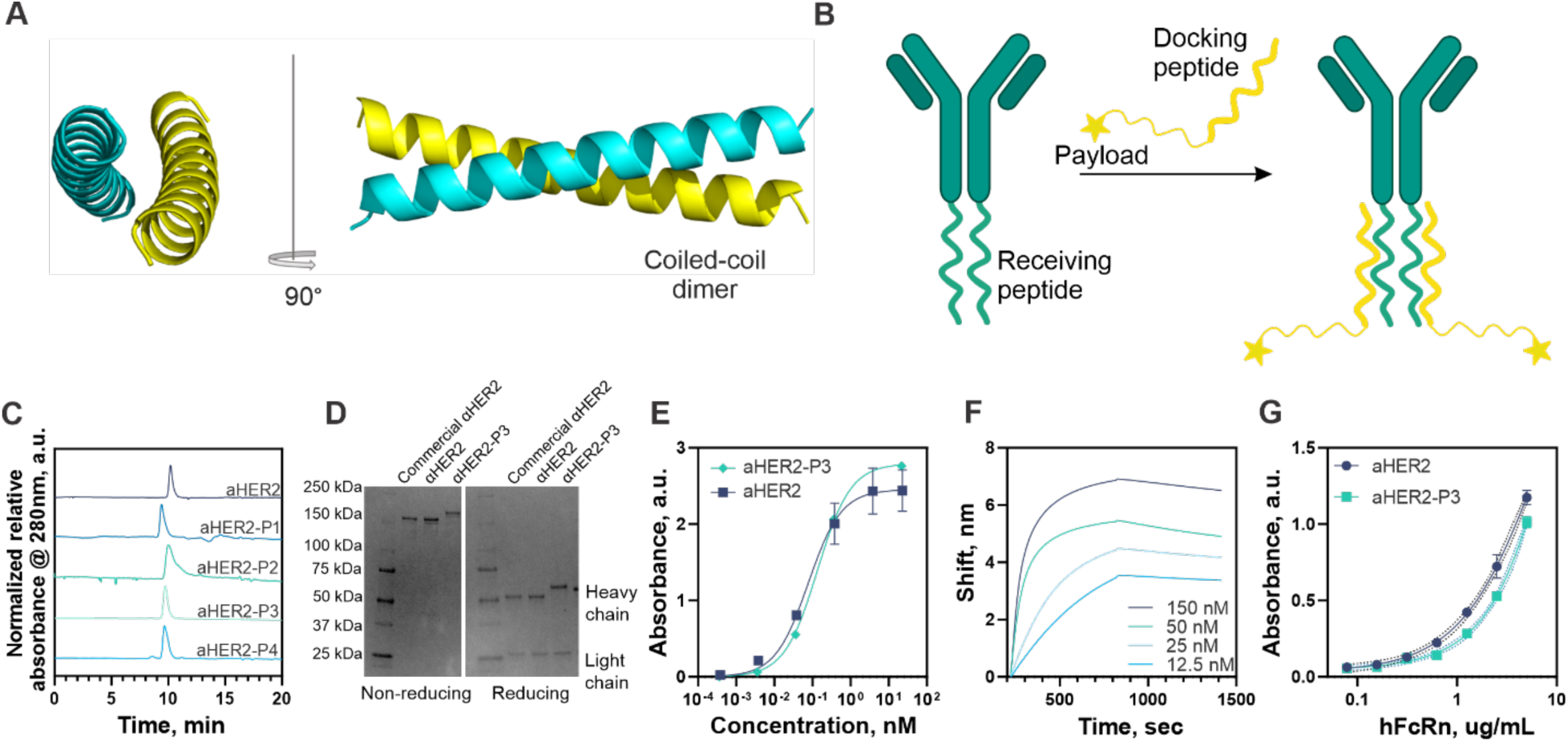
**A.** AlphaFold illustration of coiled-coil heterodimer formed from the receiving and docking peptides **B.** Schematic diagram illustrating drug loading process to an antibody through formation of coiled-coil structure between docking peptide conjugated to the drug and receiving peptide fused to the heavy chain of an antibody **C.** SEC-UV traces at 280 nm of native trastuzumab (αHER2) and trastuzumab fused to different receiving peptides (αHER2-P1, αHER2-P2, αHER2-P3, αHER2-P4) **D.** Coomassie-stained SDS-PAGE gel analysis under non-reducing (left) and reducing (right) conditions of commercial trastuzumab (commercial αHER2), in house produced native trastuzumab (αHER2) and trastuzumab fused to P3 peptide (αHER2-P3) **E.** ELISA assay demonstrating antigen-binding ability of native αHER2 and αHER2 fused to P3 peptide (αHER2-P3) **F.** Binding affinity of αHER2-P3 to biotinylated P4 peptide measured by biolayer interferometry (BLI)**. G.** ELISA measurements of binding of native αHER2 and peptide modified αHER2-P3 antibodies to human FcRn receptor. Data are presented as mean ± s.d., n = 3.

Here, we report a novel biorthogonal platform based on supramolecular assembly of coiled-coil peptides for preparing novel antibody-conjugates, including antibody-drug conjugates (ADCs). The Fc region of an IgG with two heavy chains is poised for a heterodimer coiled-coil supramolecular interaction: two C-terminal receiving peptides on the antibody assemble with two separately synthesized docking peptides containing the conjugated payload of interest (**Figure 1B**). This facile synthetic strategy enables uniform stoichiometric site-specific antibody conjugation while maintaining antibody functionality. It is a convergent bioconjugation method where the final self-assembly step occurs under mild aqueous conditions facilitating an easy, modular pairing of antibodies to payload(s) simplifying final production and purification. We site-specifically pair antibodies with fluorophores, cytotoxic drugs, peptides, polymers, nucleic acids, and enzymes. The resulting antibody-conjugates retain function and bioactivity, and a prepared ADC excels in a murine xenograft human ovarian cancer model.

## RESULTS

### Antibody-receiving peptide production and characterization

We identified two sets of peptides which form dimeric coiled-coil structures without self-interaction to create the bioconjugate platform (47). Each of the four peptide sequences (P1, P2, P3 and P4) comprise four amino acid heptad repeats containing different patterns of glutamic acid or lysine for promoting electrostatic interaction between the coils and leucine or isoleucine for hydrophobic interaction (**SI Table 1**). We introduced a single unnatural azide-bearing amino acid into the C-terminus of these peptides to enable subsequent conjugation, generating a novel set labeled as - P1N_3_, P2N_3_, P3N_3_, P4N_3_. Small peptides bearing unnatural amino acids are readily synthesized via solid-phase peptide synthesis at milligram scale (48,49) unlike large antibodies that usually require elaborate optimization of genetic encoding and protein expression (11,50,51). Individually, the peptides do not form an organized structure but spontaneously form a heterodimeric coiled-coil structure when mixed at 1:1 molar ratio with its pairing peptide (P1N_3_ when mixed with P2N_3_, and P3N_3_ when mixed with P4N_3_). The assembled coiled-coil structure is stable over time and under endosomal-lysosomal pH conditions (**Figure SI1**).

Next, we designed a set of plasmids encoding the trastuzumab heavy chain fused at the C-terminal end with one of the receiving peptide sequences (P1, P2, P3, or P4; **Figure SI2**). We expressed the antibodies in Expi293F cells and collected the protein via Protein A column chromatography. A fully folded antibody possesses two identical peptide tags linked to the two heavy chains at the Fc region. The far-UV CD spectrum displays a broad minimum at 218 nm for the unmodified αHER2 antibodies and for the peptide-tagged antibodies (αHER2-P1, αHER2-P2, αHER2-P3, and αHER2-P4), characteristic of a significant presence of β-sheets in IgG (**Figure SI3**) indicating that coil tag does not interfere with the protein folding. The peptide-modified antibodies elute slightly earlier (αHER2-P1 t=9.2 min; αHER2-P2 t=10.0 min; αHER2-P3 t=9.2 min; αHER2-P4 t=9.2 min) than the native αHER2 antibodies (t=10.1 min) during SEC chromatography, consistent with an increase in mass due to peptide insertion (**Figure 1C**). Antibodies modified with the P3 peptide display a more uniform peak, and, thus, we used this antibody for further experiments. SDS-PAGE analysis additionally confirms the modification of the heavy chain with the coil peptide (**Figure 1D**). Non-reduced antibodies with P3 inserted peptide exhibit a shift toward the higher molecular weight region compared to unmodified antibodies. Upon reduction, the light chain remains intact, while the heavy chain -P3 conjugates shifts to the higher molecular weight region, as anticipated based on the conjugate design. Finally, the MALDI-TOF MS spectrum reveals a major peak at 156 kDa for αHER2-P3, corresponding to the expected molecular weight of the full antibody (trastuzumab M_w_ ≈ 148 kDa) with two receiving peptides (P3 M_w_ ≈ 4 kDa each) (**Figure SI4**).

To confirm the recombinant antibody bind the antigen (ErbB2/Her2 protein), we performed an ELISA assay and compared it to the commercial *α*HER2 without the peptide tag and *α*HER2 with a P3 receiving peptide tag. The EC_50_ values for all tested samples are: 0.110 ± 0.024 nM and 0.133 ± 0.003 nM respectively, indicating that the peptide tag does not compromise target binding of the antibody (**Figure 1E**).

We then measured the affinity of coiled-coil formation when one of the peptides is fused to the antibody. We immobilized the biotin-labeled docking peptide (P4N_3_-biotin) on a BLI sensor and exposed it to αHER2-P3 receiving peptide antibody at various concentrations. A shift in the interference pattern occurs only between the complementary pair of a docking peptide (P4N_3_) and the corresponding P3 receiving peptide on αHER2 antibodies (αHER2-P3). The calculated dissociation constant (Kd) is 6 x 10^-10^ M, indicating a highly stable structure (**Figure 1F**, **SI5**). The interaction is specific as antibodies without a peptide tail and those with a non-specific peptide tail, control groups, do not elicit a BLI response.

Next, we evaluated if conjugation at the Fc region interferes with its interaction with the neonatal Fc receptor (FcRn), which regulates IgG recycling. We plated native trastuzumab antibody or αHER2-P3 on an ELISA plate and incubated with biotin-labeled human FcRn receptor, followed by detection with streptavidin-HRP conjugates. The αHER2-P3 antibodies maintain their affinity to FcRn after introducing the coil, however the binding is slightly lower than trastuzumab (**Figure 1G**). Overall, these findings indicate that the introduction of a coiled-coil peptide tag preserves the key properties of the mAb and its antigen-binding capabilities.

### Conjugation principle

With introduction of a coil peptide tag preserving the properties of the mAb and its antigen-binding capabilities, we next investigated payload loading onto the antibody which occurs in two steps: 1) attachment of the payload to the docking peptide via a strain-promoted azide-alkyne cycloaddition reaction; and 2) self-assembly of the payload-conjugated docking peptide to the receiving peptide on the antibody. This convergent approach enables expanded payload chemistry on the docking peptide independent of antibody modification, thereby separating the steps, minimizing the reactions with the antibody, and avoiding denaturation or aggregation of monoclonal antibody in organic solvents and/or harsh pH conditions. Thus, this facile plug-and-play supramolecular assembly opens opportunities for novel chemistries and payloads to be conjugated to the docking peptide.

### Docking peptide conjugation

To demonstrate the utility of the methodology, we prepared docking peptides linked with different entity classes to include a fluorophore (DBCO BP Fluor 647), antimitotic agents mertansine (DM1) and monomethyl auristatin E (MMAE), a DNA oligo (polyACCC), a polymer (e.g., PEG_10k_), an enzyme (e.g., HRP), a biotin molecule, and a phospholipid (DSPE). DBCO modified payloads, except HRP, were mixed is a small excess (1.2 eq) with azide bearing docking peptide (P4N_3_) (**Figure 2A**). The coupling reactions occur in minutes as monitored through depletion of DBCO absorbance peak at 310 nm (**Figure 2B**). The estimated reactions half-life with P4N_3_ is 26 ± 2 min for DBCO BP Fluor 647, 12 ± 3 min for DBCO-PEG_4_-DM1, 42 ± 10 min for DBCO-PEG_4_-VC-MMAE, 77 ± 13 min for polyA_15_CCC oligo, 27 ± 4 min DBCO-PEG_10k_, 16 ± 2 min for DBCO-PEG_4_-biotin, and 25 ± 8 min DBCO-PEG_4_-DSPE (**Figure 2C-I**). Mass spectrometry confirms the sequential conjugation and desired product formation (**Figure SI6**).

**Figure 2.**
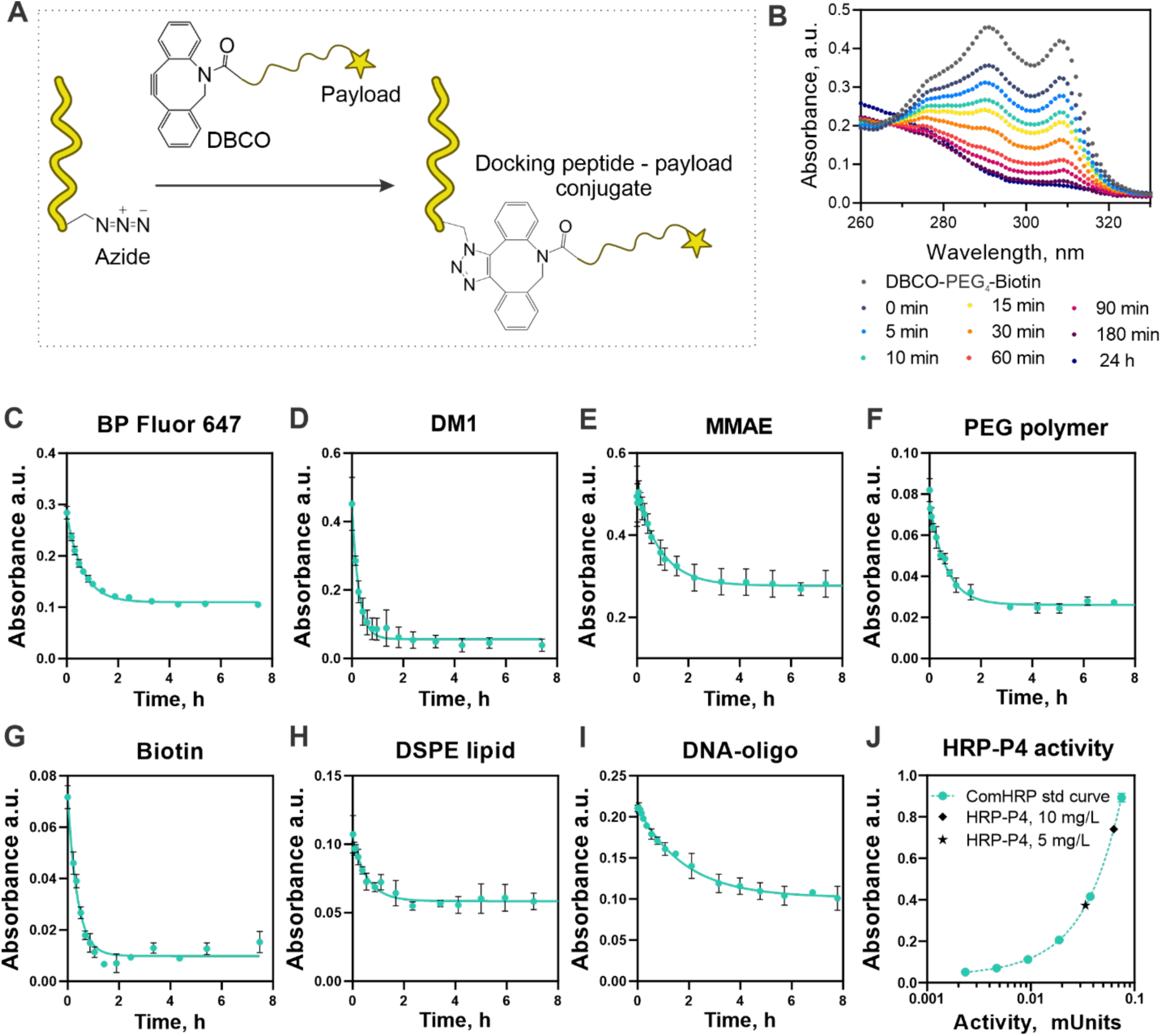
**A.** Schematic illustration of azide-modified coil peptide reacting with DBCO-bearing payload **B.** DBCO-PEG_4_-biotin absorbance spectra change upon conjugation to the P4N_3_. At the beginning of reaction (0 min) the intact DBCO-moiety absorbs in the UV spectrum with a characteristic maxima at 310 nm and stops absorbing when the structure is rearranged as a result of click conjugation **C-I**. Conjugation kinetics of DBCO-modified payloads to azide-modified docking peptide P4-N_3_ at room temperature in PBS buffer solution (pH 7.4): **C**. DBCO BP Fluor 647, **D.** DM1-PEG_4_-DBCO, **E.** MMAE-CV-PEG_4_-DBCO, **F.** DBCO-mPEG (Mw 10,000), **G.** DBCO-PEG_4_-biotin, **H.** DBCO-PEG_4_-DSPE, **I.** 5’-DBCO-N-polyA_15_GGG, **J.** TMB-based HRP activity assay measuring the activity of recombinant HRP-P4 conjugate compared to commercially available native HRP (comHRP). Data are presented as mean ± s.d., n = 3.

To enable assembly of antibody-enzyme conjugate using this technology, we expressed a recombinant enzyme with the docking peptide sequence fused into the C-terminal of the peroxidase C1A. The activity of the engineered HRP-P4 enzyme is > 60 U/mg, as measured via an enzymatic reaction with the colorimetric 3,3ʹ,5,5ʹ-tetramethylbenzidine (TMB) substrate (**Figure 2J, SI7**). The introduction of peptide to the HRP enzyme doesn’t interfere with its activity and provides a handle for assembly with receiving peptide.

### Coiled-coil mediated assembly of antibody–payload conjugates and their functions

We then assembled the antibody conjugates by simply mixing the antibody-receiving peptide (αHER2-P3) with the payload modified docking peptide in PBS for one hour followed by spin column centrifugation to isolate the desired product (**Figure 3A**). To confirm the conjugation and payload’s activity, we performed functional assays. The antibody conjugate with the BP Fluor 647 payload absorbs at 647 nm while the native antibody does not (**Figure SI7**). Based on absorbance measurements at 280 nm and 647 nm, the fluorophore-to-antibody ratio is 2.02, consistent with the designed coiled-coil assembly. The resulting antibody-fluorophore conjugate specifically targets ErbB2/HER2-positive cells in an ErbB2/Her2+ dependent manner as determined by flow cytometry using this fluorescent tag (**Figure SI8**).

**Figure 3.**
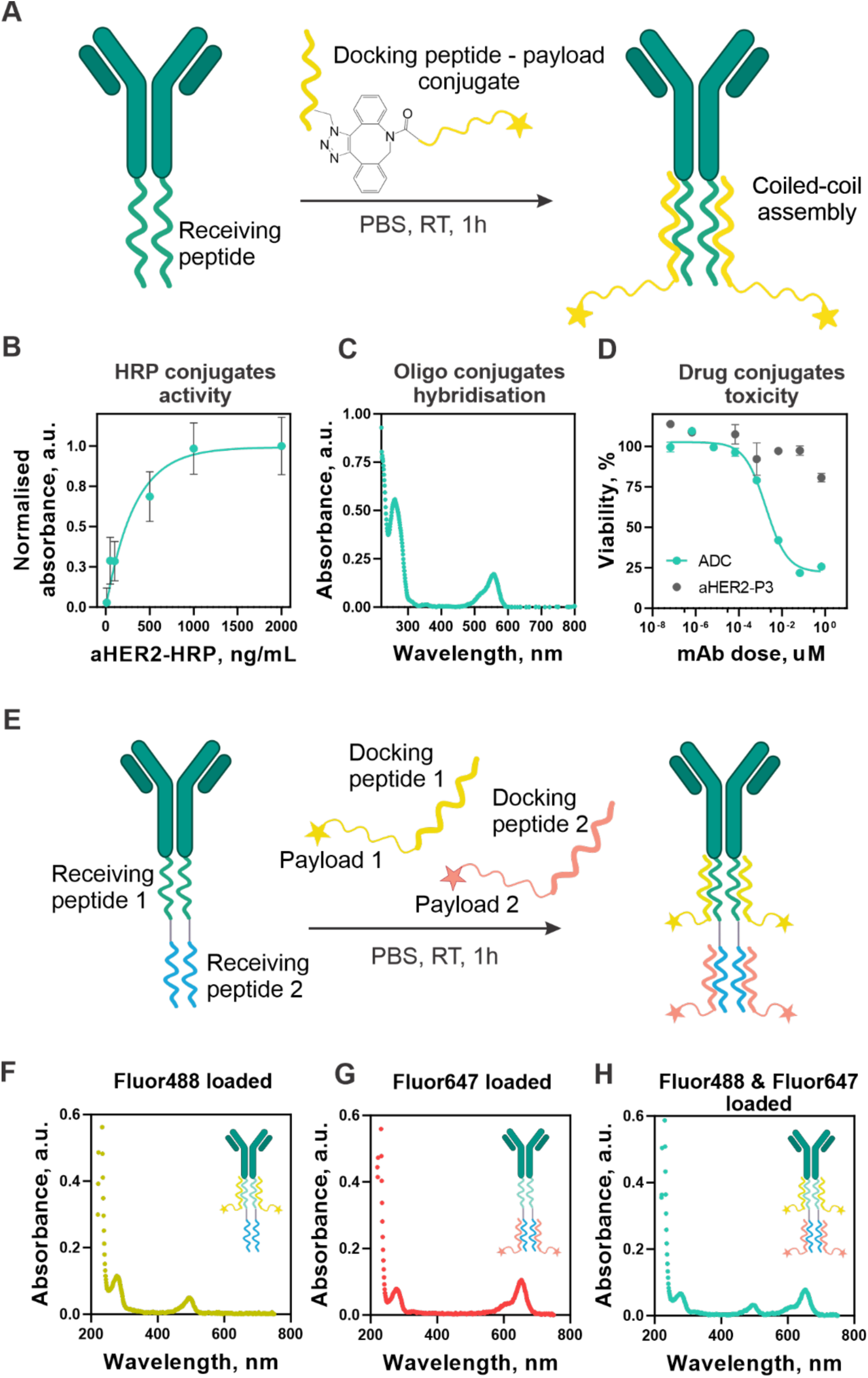
**A.** Schematic illustration of an antibody reacting with payload-modified coil peptide resulting into antibody-payload conjugate **B.** Direct ELISA assay assessing the binding and enzyme activity of αHER2-HRP conjugate **C.** UV-Vis analysis of antibody conjugated to polyA_15_GGG oligo annealed with a fluorophore labeled polyT_15_CCC-TAMRA oligo **D.** Cytotoxicity of the αHER2 ADC containing MMAE drug against SKBR-3 cells compared to equivalent unloaded αHER2-P3 antibody concentrations over 72 hours **E.** Schematic illustration of an antibody reacting with two orthogonal payload-modified coil peptides resulting into antibody bearing two different payloads **F-H.** UV-Vis analyses of antibodies assembled with single **F.** Fluor488 payload or **G.** Flour647 payload and **H.** dual-loaded conjugates.

To evaluate the αHER2-HRP conjugate activity, we conducted a direct ELISA using ErbB2/Her2 protein immobilized on a plate and TMB as the HRP substrate (**Figure 3B**). The αHER2-HRP conjugate catalyzes the oxidation of TMB in a dose-dependent manner, demonstrating that, despite the presence of a large payload, αHER2-HRP retains its binding affinity to ErbB2/Her2 post-conjugation, while also maintaining HRP enzymatic activity. We performed indirect ELISA assays for antibody-PEG_5k_-DSPE lipid, mPEG_10k_ polymer, and biotin conjugates using anti-PEG secondary antibodies or streptavidin-HRP for payload detection. All the assays confirm the successful conjugation of antibodies with different payloads (**Figure SI9**). For oligonucleotide-loaded antibodies, we mixed the single-stranded DNA-loaded (polyA_15_GGG) antibody with a complementary TAMRA fluorophore-labeled strand (polyT_15_CCC-TAMRA) and incubated the mixture to allow hybridization followed by purification via spin filtration. The results confirmed successful hybridization on the complementary oligos, as indicated by the TAMRA fluorophore absorbance of the final product, demonstrating the capability of formation of a double-stranded oligonucleotide structure post conjugation to the αHER2 antibody (**Figure 3C, SI11**). Finally, we assessed the cytotoxicity of the synthesized αHER2-MMAE conjugates using the MTS assay on three cell lines with different HER2 expression levels: 4T1 (ErbB2/Her2-), A549 (Low ErbB2/Her2+), SKOV-3 (Medium-high ErbB2/Her2++) and SKBR3 (High ErbB2/Her2+++). We treated cells with varying concentrations of ADCs, unloaded αHER2 antibodies, or free MMAE drug for 72 hours (**Figure 3D, SI12**). Treatment with the ADC shows a dose-dependent and ErbB2/Her2 expression-dependent decrease in cell viability for SKBR-3 (ADC EC_50_ ∼ 0.002 uM), SKOV-3 (ADC EC_50_ ∼ 0.09 uM) and A549 cell lines, while showing minimal toxicity in ErbB2/Her2 negative 4T1.

### Dual-payload loaded antibody-conjugates

An additional beneficial feature of this coiled-coil design is that two or more, similar or different, entities can be attached to the antibody by extending the coil-coiled structure with the same peptide or an orthogonal peptide sequence set. To demonstrate the assembly of dual-payload loaded antibodies, we fused receiving peptide P1 followed with another receiving peptide P3 sequence into the heavy chain of trastuzumab resulting in antibodies with two sequential orthogonal conjugation sites (**Figure 3E, SI13**). As a model system, we chose DBCO-modified BP Fluor 488 (Fluor488) and DBCO-modified BP Fluor 647 (Fluor647) which we conjugated to azide-modified P2N_3_ and P4N_3_ peptides respectively resulting into P2-Fluor488 and P4-Fluor647 conjugates. Coil peptide P1 forms an orthogonal coiled-coil structure with peptide P2, while peptide P3 forms a dimer with peptide P4. Next, we mixed P2-Fluor488 or P4-Fluor647 with the antibodies separately. After one hour incubation and spin-filtering purification we obtained the conjugates bearing either Fluor488 or Fluor647 fluorophores. The absorbance spectra of the conjugation products exhibit maxima at 280 nm, corresponding to protein absorbance, and either at 488 nm or 647 nm, depending on which payload loaded indicating that the extended peptide tag structure does not interfere with payload loading. (**Figure 3F,G**). We then mixed both P2-Fluor488 and P4-Fluor647 with antibodies bearing P1 and P3 peptides. The corresponding peptide pairs self-assemble leading to formation of dual-loaded antibody conjugates both with Fluor488 and Fluor647 (**Figure 3H**). This approach offers the opportunity to increase the drug-antibody ratio or load two different payloads in a defined stoichiometric ratio and at a specific location on the antibody, enabling the development of new co-delivery strategies.

### Stability of antibody-payload conjugates

To evaluate the stability of the proposed conjugation method, we compared the stability of conjugates formed via conventional thiol-maleimide covalent linkage with those generated through coiled-coil-mediated conjugation. In addition to the P3 and P4 pair (4 repeating amino acid heptad pattern, total 28-amino-acid-long coil peptides) described above, we investigated two additional peptide pair designs based on prior work on coiled-coil interactions. Woolfson et al., Xia et al., and Lee et al. report stabilized coiled-coil regions through increasing the lengths of the coiled-coil regions (34) or the presence of complementary reactive groups capable of forming stabilizing covalent bond between the peptides (44,52). For the first design, we extended the P3 and P4 peptide sequence by one coiled-coil heptad pattern, creating 35 amino acids long peptides (P3_5mer_ and P4_5mer_N_3_, see **SI Table 1**). We used the same repeating units for the second design except that we replaced the isoleucine with a cysteine residue in the last heptad unit of each peptide to allow a disulfide bond formation between coils (P3_5mer_-cys and P4_5mer_-cysN_3_, see **SI Table 1**).

We produced αHER2-P3, αHER2-P3_5mer_ and αHER2-P3_5mer_-cys antibodies (**Figure SI14**) and assembled them with corresponding docking peptides conjugated to DBCO BP Fluor 647. We then incubated the antibody-fluorophore conjugates (AFCs) in human serum at 37°C in 5% CO_2_ atmosphere. We collected serum samples over a 28-day period and analyzed for intact AFC levels using an ELISA assay that detects both the payload and antibody portion of AFC. The AFC prepared by the standard thiol-maleimide hinge conjugation technique degrades by 25%, while the coiled-coil based conjugates showed varying stability levels depending on the coil design (**Figure 4A,B**). The P3-P4N_3_ coils disassemble over the first few days, with only 13% of the AFC intact by the end of the study. In contrast, the AFCs prepared using P3_5mer_, P4_5mer_N_3_ and P3_5mer_-cys, P4_5mer_-cysN_3_ coil pairs are stable with only a 4% loss in structure over 4 weeks, outperforming the conventional conjugation approach (**Figure 4C,D**).

**Figure 4.**
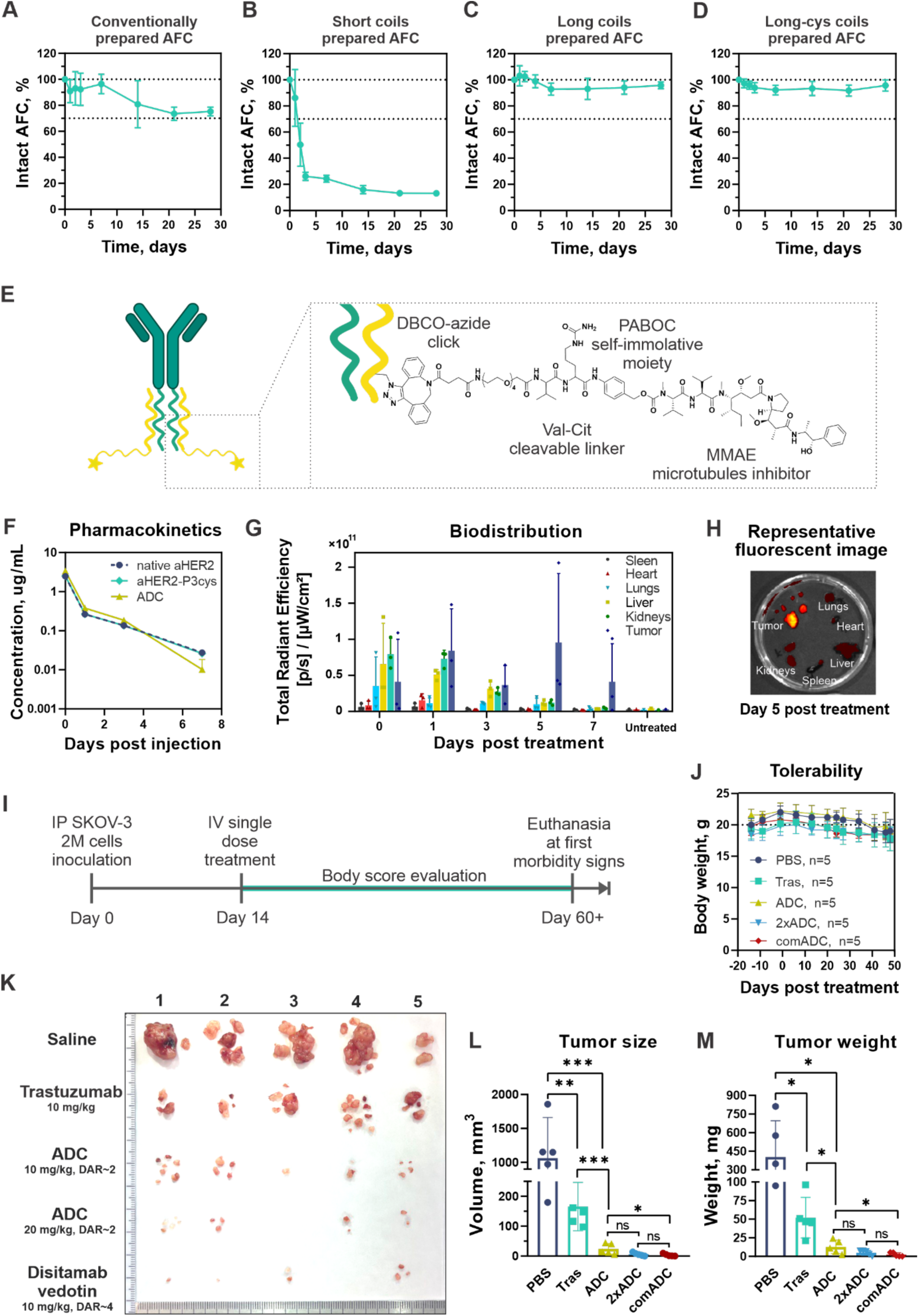
**A.** Stability of conventionally prepared AFCs via hinge cysteines conjugation, **B**. via short 28 amino acid long coiled-coil peptides, **C.** via long 35 amino acid long coiled-coil peptides and **D.** via long 35 amino acid long coiled-coil peptides with cysteine residue substitution in human plasma incubated at 37 °C. The ratio of intact AFC was determined by ELISA normalizing the detected payload amount to the total amount of IgG compared to the standard curve prepared from intact AFC (no incubation in serum, flash frozen at zero time point) **E.** Schematic illustration of ADC conjugation strategy and molecular structure of the cleavable linker-payload **F.** PK of unmodified trastuzumab (αHER2), unloaded ADC (αHER2-P3_5mer_-cys), and fully assembled coiled-coil ADC (ADC) in female Balb/c (N = 3, n=6 for trastuzumab and ADC; N = 2, n=4 for unloaded ADC group). Blood was collected at indicated time points and the total amount of human IgG was quantified by indirect ELISA using standard curve **G.** Ex vivo organ biodistribution of AFC injected i.v. at 10 mg/kg dose into SKOV-3 tumor bearing BALB/c nude mice at different time points (n=3 per time point). The total radiant efficiency was calculated using IVIS software keeping area of ROI the same across all the samples. **H**. Representative fluorescent picture of ex vivo organs post AFC injection on day 5. **I.** Orthotopic xenograft model of ErbB2/Her2-positive cancer study design and timeline. Female NSG mice were engrafted with 2 million SKOV-3 tumor cells by intraperitoneal injection, were randomized on day 14 and treated with single dose of PBS or trastuzumab antibody (10 mg/kg) or coiled-coil ADC (10 mg/kg) or high dose coiled-coil ADC (2xADC, 20 mg/kg) or commercial ADC disitamab vedotin (10 mg/kg), n= 5 mice per group. All mice were euthanized on day 48 post-treatment, when PBS-treated mice exhibited severe hunching posture and signs of morbidity, to record endpoint tumor volume and mass **J**. Average mouse body weight measured twice a week (±SD) for each treatment group demonstrating no significant difference between the groups over the observation period. **K.** Images of harvested tumors at study termination at day 48 post treatment. **L.** Recorded tumor size and **M.** tumor mass. Statistical significance is indicated as follows: ns, non-significant; *P ≤ 0.05, **P ≤ 0.01, ***P ≤ 0.005 (unpaired t-test).

### Efficacy of antibody-drug conjugates in vivo

Given our interest in ovarian cancer and the dearth of available treatments (53–56), we prepared MMAE loaded ADC conjugates to assess the overall therapeutic potential of the coiled-coil conjugation. We conjugated P4_5mer-_cysN_3_ to DBCO-PEG_4_-Val-Cit-PAB-MMAE (**Figure 4E, Figure SI15A**), where the Val-Cit-PAB portion ensures scar-less drug release (57). MMAE-peptide loading to the αHER2-P3_5mer_-cys antibody results in a SEC trace shift (**Figure SI15B**) with drug-antibody ratio of two as determined by UV-vis absorbance at 247 nm and 280 nm (58,59) (**Figure SI15C**). An indirect ELISA using anti-MMAE secondary antibodies additionally confirms the modification of αHER2-P3_5mer_-cys with MMAE drug compared to unloaded antibodies (**Figure SI16**).

First, we evaluated the pharmacokinetic (PK) profile of the ADC *in vivo*. We intravenously (i.v.) administrated native mAb (trastuzumab), unloaded αHER2-P3cys mAb, and MMAE loaded ADC at dose of 10 mg/kg to healthy Balb/c mice. We collected blood samples at predefined time points and measured the concentration of mAb in the blood by an ELISA assay. The half-life for trastuzumab is 6.5 hours (95% CI 5.5-7.3 h), similar to those of the unloaded ADC and MMAE loaded ADC conjugates at 6.8 (95% CI 5.0-7.7 h) and 6.5 (95% CI 6.2-7.4 h) hours respectively (**Figure 4F**). The coiled-coil peptide tag does not negatively impact antibody PK and clearance. These data along with the *in vitro* results showing that the antibody binds the Fc receptor (**Figure 1G**) suggests that coiled-coil peptide modification of the C-terminus does not significantly affect FcRn binding.

To assess tissue biodistribution of the antibody conjugate, we used an orthotopic xenograft model of ErbB2/Her2-positive human ovarian tumors by implanting SKOV-3 cells intraperitoneally into immunodeficient BALB/c nude mice. Then tumors established, we i.v. administered a BP Fluor 647-loaded AFC at a dose of 10 mg/kg. At 2 hours, 1 day, 3 days, 5 days, and 7 days post-injection, we euthanized the animals and quantified the fluorescent signal in the tumor and major organs by IVIS. (**Figure 4G,H, Figure SI17**). The AFC accumulates in the tumor throughout the 7-day study period, along with liver kidneys and lungs at the earlier time points. These results are consistent with the literature (60,61) and demonstrate the ability to enhance accumulation of payload to ErbB2/Her2-positive tumors.

Using the established orthotopic xenograft model (**Figure 4I**), we performed an efficacy study and i.v. administered a single dose of 10 mg/kg of trastuzumab (native αHER2 antibodies) or 10 mg/kg of coiled-coil ADC (n = 5 per group). Additionally, we administered PBS as the negative control and disitamab vedotin — the latest HER2-targeting ADC to advance to Phase III clinical trials (62) — as the positive control. It consists of the hertuzumab antibody and, similar to our ADC, uses the cytotoxic payload MMAE linked via a Val-Cit cleavable linker. However, disitamab vedotin employs cysteine-based conjugation, resulting in approximately four MMAE molecules per antibody. To match the drug dose of positive control, we also administered a doubled dose of coiled-coil ADC (20 mg/kg, n = 5 per group). All groups tolerated the treatment with no significant changes in body mass (weight loss of >15%) (**Figure 4J**) or observable adverse effects on animal behavior at earlier time points. However, at week 7, the PBS treatment group and one trastuzumab-treated animals exhibited hunched posture and signs of gait rigidity. We euthanized the mice on day 48 post single dose treatment, and tumor size and mass were evaluated. Compared to the PBS treated animals, all treatment groups significantly reduce tumor size and mass (**Figure 4K-M**). Mice treated with 10 mg/kg coiled-coil ADC exhibit a significant reduction in tumor size, with a 31-fold decrease in tumor mass and 44-fold decrease in tumor volume (m = 13±9 mg, V = 24±18 mm^3^) compared to the untreated control (m = 403±290 mg, V = 1062±599 mm^3^). In contrast, those receiving the antibody vehicle control (trastuzumab) show only an 8-fold and 7-fold reduction, respectively (m = 52±27 mg, V = 166±81 mm^3^). Treatment with 20 mg/kg ADC or disitamab vedotin completely eradicates tumor in one mouse from each group, with average tumor masses of 5±3 mg and 2±3 mg, respectively, with a similar trend in the tumor volume: V = 6±5 mm^3^ and V = 4±4 mm^3^,respectively. Disitamab vedotin is highly efficacious, consistent with its elevated drug-to-antibody ratio and enhanced affinity for the HER2 target relative to trastuzumab (63). Our comparable results from coiled-coil ADC treatment group bodes well for further use of the coiled-coil conjugation method for antibody conjugation.

Finally, we confirmed the dependency of the therapeutic efficacy of the ADC on ErbB2/Her2 expression levels *in vivo*. We compared ErbB2/HER2 expression in human lung A549 and SKOV-3 cell lines, as well as in their corresponding tumor tissues. SKOV-3 cells exhibit an 83-fold higher ErbB2/Her2 targeting at the cellular level compared to isotype control and an 11-fold increase for tumor tissues, whereas for A549 both for the cell line and tumor tissue the ErbB2/Her2 targeting is modest, with 1.6-fold and 2.5-fold increases, respectively (**Figure SI18**). We then established an A549 xenograft model after SC implantation. At the 10 mg/kg i.v. dose, neither the trastuzumab antibody (mAb), MMAE coiled-coil ADC, nor disitamab vedotin demonstrate any significant tumor growth inhibition, consistent with the low ErbB2/Her2 expression levels observed in A549 cells and tumors (**Figure SI19**).

## CONCLUSION

We describe a facile method for site-specific antibody conjugation that enables uniform loading of distinct payloads under mild aqueous conditions. Coiled-coil conjugation perseveres both antibody antigen binding sites for target engagement and heavy chains constant domains for Fc binding and recycling. The convergent approach bifurcates the conventional antibody method thereby minimizing the reaction steps and conditions exposed to the antibody while increasing the synthetic flexibility for payload conjugation as it employs peptide-based chemistry. In doing so, this strategy also provides access to readily synthesizing several antibody conjugates or a library of antibody conjugates whereby one or more antibodies are prepared followed by mixing and matching with different conjugates. Additionally, the step-wise assembly of the coiled coils affords relative spatial control over the site-specific location of multiple conjugates on the antibody. The coiled-coil composition controls conjugate stability and structures with stability beyond that of conventional hinge cysteine conjugation which is known to prematurely release payload. An MMAE-ADC targeting ErbB2/Her2+ tumors, prepared via self-assembly, significantly reduces tumor volume in a human ovarian cancer xenograft model outperforming the antibody alone with validated similar performance against best-in-class therapeutics. Heterodimer coiled-coil driven bioconjugation opens new avenues for well-defined stoichiometric site-specific antibody labeling with one or more payload compositions and as a supramolecular chemistry approach expands existing biorthogonal strategies.

## MATERIALS AND METHODS

### Cells

SKOV-3, SKBR-3, A549 and 4T1 cell lines were obtained from American Type Culture Collection (ATCC, USA). Cell media supplemented with 10% fetal bovine serum (FBS) and 1% penicillin/streptomycin was used to maintain the cell lines: McCoy’s 5A Medium (ATCC 30-2007) for SKBR-3 and SKOV-3, F-12K Medium (ATCC 30-2004) for A549 and RPMI 1640 (ATCC 30-2001) for 4T1. The cell lines were cultured at 37 °C in a humidified CO_2_ (5%) incubator. Expi-293F cells (Gibco™) were cultured in Expi293™ Expression Medium (Gibco™) at 37 °C under constant shaking (120 r.p.m.) in a humidified CO_2_ (8%) incubator.

### Plasmids

The pcDNA3.1(-) backbone was pre-cut with EcoRV and BamHI restriction enzymes (NEB) and purified using the QIAquick PCR Purification Kit (Qiagen). The 5ʹ ends were dephosphorylated with Quick CIP (NEB) to prevent re-ligation. The gBlocks of interest were synthesized by IDT and inserted into the prepared pcDNA3.1(-) backbone via Gibson Assembly (NEB). The assembled plasmids were transformed into NEB Stable *Escherichia coli* and purified using the ZymoPURE II Plasmid Midiprep Kit (Zymo Research). Sequence confirmation was performed by Sanger sequencing using CMV-F and BGH-R primers (Genewiz).

### Protein expression and purification

Native αHER2 and receiving-peptide modified αHER2 antibodies, as well as HRP-P4 conjugates were produced in Expi293F expression system (Gibco™) according to manufactures’ protocol. Expi293F cells were co-transfected with a mixture of light and heavy chain containing plasmids for mAb production or with a single plasmid containing HRP-docking peptide fusion sequence. Proteins were harvested on day 6 by centrifugation at 3,220 g for 20 min. The supernatant was filtered through a 0.22-µm membrane for subsequent purification. mAbs were purified using a gravity-flow column packed with rProtein A Sepharose Fast Flow resign (Cytiva), HRP-P4 using HisPur™ Ni-NTA Resin (Thermo Fisher Scientific). Collected proteins were concentrated and buffer exchanged with centrifugal filters (10kDa or 50 kDa molecular weight cut-off, MilliporeSigma). The concentration of all proteins was determined by absorbance at 280 nm using sequence predicted extinction coefficients (mAb ε_280_=216,340 M^-1^cm^-1^; and HRP-P4 ε_280_=15,065 M^-1^cm^-1^).

### Proteins characterization

#### Size exclusion chromatography (SEC)

Antibodies were analyzed using an AdvanceBio SEC column (300Å, 2.7 µm, 4.6 x 150 mm Agilent Technologies). The column was equilibrated and operated with phosphate-buffered saline (PBS, pH 7.4) as the mobile phase at room temperature under isocratic conditions at 0.8 mL/min flow rate, and elution was monitored by UV absorbance at 280 nm (Varian ProStar HPLC).

#### SDS-PAGE gel

Produced proteins (αHER2, αHER2-P3, HRP-P4) were separated by SDS polyacrylamide gel electrophoresis (SDS-PAGE). 1 μg of each protein was mixed with gel loading buffer (reducing or non-reducing) and loaded into wells of a 4-15% precast gel (BioRad). The gel was subjected to 120 constant volts for 40-60 minutes. Following this, the gel was stained with Coomassie Blue (Thermo Fisher Scientific).

#### Circular Dichroism (CD)

Antibodies and peptides were prepared in PBS buffer (pH 7.4) at the concentration of 0.2 g/L. The measurements were performed on CS/2 Chirascan (Applied Photophysics) CD spectrometer using a 1 mm QS 100 (Hellma) cuvette. PBS solution was used as a blank. To study the stability of the coiled-coil assembly at different pH levels, the peptides were buffer exchanged using 3 kDa MWCO spin filters (MilliporeSigma) into a PBS solution adjusted with HCl to pH 5.4, 6.4, or 7.4. For the overtime stability study of the P3-P4 coiled-coil assembly, the peptides were incubated at 37 °C between measurements. The proportion of intact coiled-coil assemblies relative to single α-helices was estimated by calculating the ratio of ellipticity at 222 nm to that at 208 nm (64).

#### Recombinant antibodies antigen binding assay

The ability of recombinant antibodies to recognize ErbB2/Her2 protein was evaluated using indirect ELISA. Nunc MaxiSorp 96-well ELISA plates were coated overnight at 4 °C with 1 μg/mL of recombinant human ErbB2/Her2 protein (Cat. #10126-ER, R&D Systems Inc.) in 0.05 M carbonate–bicarbonate buffer (pH 9.6). Plates were then blocked with 5% bovine serum albumin (BSA) in PBS and washed with PBS containing 0.05% Tween-20 (PBS-T) between each step, following a standard ELISA protocol. Serial dilutions of native αHER2 (Human ErbB2/Her2 Antibody, Cat. # MAB9589-100, R&D Systems Inc) or antibodies fused to receiving peptide αHER2-P3 were prepared in blocking buffer (starting concentration 3 ug/mL). Wells without antigen were used as negative controls. Goat anti-human IgG H&L (Abcam) was used as the secondary antibody. After washing, 100 μL of TMB substrate (1-Step™ TMB ELISA Substrate Solution, Thermo Scientific™) was added to each well. Following incubation, the reaction was stopped with 100 μL of 2N sulfuric acid. Absorbance was measured at 450 nm using a microplate reader (SpectraMax iD3, Molecular Devices).

#### *In vitro* FcRn recognition assay

The interaction between recombinant antibodies with fused receiving peptides was studies according to the SinoBiological activity protocol for recombinant human FCGRT & B2M heterodimer protein (Cat. # CT071-H27H-B, SinoBiological). Nunc MaxiSorp 96-well ELISA plates were coated overnight at 4 °C with 2 μg/mL of native αHER2 or antibodies fused to receiving peptide αHER2-P3 in 0.05 M carbonate–bicarbonate buffer (pH 9.6). Plates were then blocked with 5% bovine serum albumin (BSA) in PBS and washed with PBS containing 0.05% Tween-20 (PBS-T) between each step. Biotin labeled recombinant Human FCGRT & B2M Heterodimer (Cat. # CT071-H27H-B, SinoBiological) was prepared as serial 2x dilutions (starting concentration 5 ug/mL) in dilution buffer (PBS, pH=6.0, with 0.2% Tween-20, 0.1%BSA) and incubated in ELISA plate for 1 hour at 37 °C. A solution of streptavidin/HRP in Dilution buffer (0.5% BSA in PBST, pH=6.0) was used for detection. After washing steps, 60 μL TMB solution (1-Step™ TMB ELISA Substrate Solutions, Thermo Scientific™) was added to the wells. After incubation, the reaction was stopped with 60 μL of 2N sulfuric acid. Absorbance was measured at 450 nm using a microplate reader (SpectraMax iD3, Molecular devices).

#### Biolayer Interferometry (BLI)

P2N_3_, P3N_3_ and P4N_3_ peptides were first biotinylated with DBCO-PEG_4_-biotin. Streptavidin SA tips were baslined in BLI buffer (28 mM Tris-HCl pH 8.0, 5 mM MgCl, 4.25% Glycerol, 25 mM NaCl, 1.67 mg/mL BSA) and dipped into 100 nM docking peptide solution for 60 s. Then the peptide coated tips were baselined in BLI buffer again and dipped into wells containing αHER2-P3 (12.5 nM, 25 nM, 50 nM and 150 nM) or native αHER2 (100 nM) or αHER2-P2 (100 nM). After reaching the equilibrium tips were dipped in BLI buffer for complex dissociation. The assay was performed using ForteBio OctetRed96 and data analysis was performed in Octet Analysis Studio 13.0. Tips with no docking peptide loading were used as association control and tips with no dipping to antibody solutions were used as dissociation control. Trace Y-Axes were aligned to the last 5 s of the second baseline step and data was fit to a Mass Transport model.

#### Docking peptide synthesis

Docking peptides were synthesized by ABclonal (Woburn, MA) using solid phase synthesis with >95 % purity and verified by HPLC and mass spectrometry by the manufacturer.

#### Payload loading to docking peptide

The DBCO bearing payloads were dissolved according to manufacturer recommendations (BP Fluor 647 DBCO in DMF, BroadPharm; DM1-PEG_4_-DBCO in DMF, BroadPharm; DBCO-PEG_4_-VC-MMAE in DMF, BroadPharm; DBCO-NSH-polyA_15_GGG in water, Integrated DNA Technologies; DBCO-mPEG_10k_ and DBCO-PEG_2k_-OH in water, Biopharma PEG; DBCO-biotin in DMF, BroadPharm; DBCO-PEG_4_-DSPE and DBCO-PEG_5k_-DSPE in DMF:H_2_O:EtOH 1:1:2 BroadPharm). The small excess of DBCO bearing payload (1.2 eq.) was added dropwise to a solution of P4 docking peptide in PBS (20 ug). The absorbance at 310 nm was measured immediately using NanoDrop spectrophotometer.

#### MALDI-TOF analysis of peptides and antibodies

Antibodies, peptides, and their conjugates were analyzed using a Bruker Autoflex Speed MALDI-TOF mass spectrometer with sinapinic acid (SA) or CHCA (α-Cyano-4-hydroxycinnamic acid) as the matrix (10 mg/mL in 50% acetonitrile and 0.1% trifluoroacetic acid in water). Prior to analysis, samples were buffer-exchanged into ultrapure water using centrifugal filter units (50 kDa MWCO for antibodies and 3.5 kDa MWCO for peptides; Millipore). The resulting solutions were mixed 1:1 (v/v) with the SA or CHCA matrix solution immediately before spotting onto the MALDI target plate.

#### HRP-P4 docking peptide activity

Serial 10-fold dilutions of commercially available HRP (Sigma, CAS #9003-99-0) were prepared in phosphate-buffered saline (in PBS; 1 mg/L - 0.001 ug/L) as standards. A volume of 60 µL from each dilution or recombinant HRP-P4 was transferred into a well of a flat-bottom 96-well microplate. Subsequently, 60 µL of 3,3ʹ,5,5ʹ-tetramethylbenzidine (TMB) solution (1-Step™ TMB ELISA Substrate Solutions, Thermo Scientific™) was added to each well and absorbance at 650 nm was measured immediately using a microplate reader (SpectraMax iD3, Molecular devices) at one-minute intervals. In 15 minutes the rection was stopped by addition 60 uL of 2N sulfuric acid and absorbance at 450 nm was recorded. Recombinant HRP-P4 activity was quantified by comparing the end point values of recombinant HRP samples to a standard curve generated using serial dilutions of a commercially available HRP standard.

#### Antibody-payload assembly

Solution of αHER2-docking peptide (1 eq., in PBS) was mixed with corresponding payload-modified peptide (2.2 eq., in PBS) and incubated for 2 hours at RT or overnight at 4°C to let the coiled-coil peptides to self-assembly. The solution was centrifuged using a spin filter to isolate the conjugated antibody-payload.

### Conjugates functional assays

#### HRP conjugates functional assay

Activity of the αHER2-HRP conjugates was assessed by direct ELISA. Nunc MaxiSorp 96-well ELISA plates were coated overnight at 4 °C with 2 μg/mL of recombinant human ErbB2/Her2 protein (Cat. #10126-ER, R&D Systems Inc.) in 0.05 M carbonate–bicarbonate buffer (pH 9.6). Plates were then blocked with 5% bovine serum albumin (BSA) in PBS and washed with PBS containing 0.05% Tween-20 (PBS-T) between each step, following a standard ELISA protocol. The αHER2-HRP conjugates were serially diluted 2-fold in blocking buffer (2 μg/mL - 1 ng/mL, conjugates concentration determined via nanodrop using ε_280_=237,385 M^-1^cm^-1^) and incubated in ELISA plate for 1 hour at 37 °C. After washing steps, 60 μL TMB solution (1-Step™ TMB ELISA Substrate Solutions, Thermo Scientific™) was added to the wells. After 30 minutes of incubation, the reaction was stopped with 60 μL of 2N sulfuric acid. Absorbance was measured at 450 nm using a microplate reader (SpectraMax iD3, Molecular devices).

#### DNA oligo conjugates functional assay

A solution of antibodies conjugated to 5’-DBCO-NSH-polyA_15_GGG-3’ DNA oligo (αHER2-P3-P4-polyA_15_CCC, 1 eq., 0.17 mM, 27 uL in PBS) was mixed with the complementary 5’-TAMRA-CCCpolyT_15_-3’ (2.1 eq., 0.37 mM, 13 uL in water). The mixture was incubated at 34.4 °C for 30 minutes in a thermal cycler (T100 Thermal Cycler, Bio-Rad) to allow oligo hybridization. The product was purified by sequential washing using an Amicon Ultra 30 kDa MWCO spin column until no TAMRA absorbance is detected in the flowthrough (3 times). αHER2-P3 with no P4-polyA_15_CCC were used as a negative control. The absorbance spectra of the resulting conjugate were measured using NanoDrop spectrophotometer.

#### MMAE conjugates functional assay

Activity of the αHER2-MMAE conjugates was assessed by MTS cell viability assay. SKBR-3 cells were plated at a density of 5,000 cells per well, A549 cells were plated at a density of 2,500 cells per well, SKOV-3 cells were plated at a density of 2,000 cells per well, and the 4T1 were plated at a density of 1,000 cells per well the day before treatment. Next day, cell media was removed, and the cells were treated with varying concentrations of mAb (αHER2-P3), ADC (αHER2-MMAE) and free MMAE dissolved in the cell media. The MMAE was serially diluted starting from 2.5 uM in media (100 uL per well) and the ADC concentration was selected was to match the MMAE dose (2 drugs per antibody). mAb concentration was fixed to ADC concentration. After 72 hours cell media was removed and solution of MTS was added in serum free media. The signal development (30-60 min) was monitored with microplate reader (SpectraMax iD3, Molecular devices) as absorbance at 490 nm. Media only and untreated cells were used as controls.

#### Conjugates plasma stability

AFCs were prepared using both conventional hinge cysteine conjugation and coiled-coil–based strategy for subsequent stability studies. Hinge cysteine conjugation was performed via TCEP reduction (2.75 eq.). The reduction was carried out in 1 M sodium borate buffer containing 25 mM NaCl and 1 mM EDTA for 90 minutes at 37 °C. Without purification the antibodies were cooled to 0 °C and reacted with 5 equivalents of BP Fluor 647 maleimide overnight at 4 °C covered with foil. The resulting conjugates were purified using a centrifugal spin filter with a 10 kDa MWCO. Then, the conjugates were reoxidized using 2 eq. of DTNB for 20 minutes and subsequently re-purified using a 50 kDa MWCO spin filter to obtain the final product. Coiled-coil AFC were prepared as described previously using short coiled-coil peptides pairs P3-P4 or long coiled-coil peptides P3_5mer_-P4_5mer_ or cysteine modified P3_5mer_cys-P4_5mer_cys. The AFCs (5 µg) were dissolved in 0.5 mL of human plasma and incubated at 37 °C under constant shaking conditions. The aliquots samples (10 µL each) were flash frozen in liquid nitrogen and stored −80 °C for downstream analysis. The samples were analyzed using indirect ELISA assay using 2 μg/mL of recombinant human ErbB2/Her2 protein (Cat. #10126-ER, R&D Systems Inc.) as the antigen and anti-human or anti-AF647 antibodies secondary antibodies. The fluorophore detection signal was normalized to the total antibody content, as determined by absorbance using anti-human secondary antibodies. Flash-frozen AFCs collected prior to plasma incubation (day 0) were used as an intact AFC standard.

#### Animal studies

All animal experiments described in this study were performed in accordance with protocols that were reviewed and approved by the Institutional Animal Care and Use Committee of Boston University. All mice were maintained with a 12-h light/dark cycle at 20– 26 °C and 30–70% humidity in facilities accredited by the Association for the Assessment and Accreditation of Laboratory Animal Care.

#### Pharmacokinetic studies and serum preparation

Non-tumor bearing Balb/c mice (6-7 weeks old) were treated with 10 mg/kg of native mAb (αHER2) or unloaded ADC (aHER3-P3_5mer_cys) or MMAE conjugated ADC (aHER3-P3_5mer_cys -P4_5mer_cys-MMAE) intravenously (100 µL, PBS solution). Blood was collected post-treatment via tail vein bleeding at 30 minutes, 1 day, and 3 days, and via cheek bleeding at the 7-day time point. Serum was isolated by centrifuging blood in a benchtop centrifuge at 1,000 g for 10 min at room temperature. Serum was then collected, transferred to a new Eppendorf tube and stored at −20 °C for downstream analysis. The data was analyzed using indirect ELISA assay detecting total amount of human IgG and standard curve.

#### Efficacy study

Female NSG mice (6–8 weeks old) were obtained from the Jackson Laboratory (#005557). The orthotopic xenograft model of SKOV-3 was established by injecting SKOV-3 cells (2 × 10^6^ cells in 0.2 ml PBS) intraperitoneal. Mice were administered a single dose of 100 µl of PBS or trastuzumab or ADCs at a dose of 10 mg/kg or 20 mg/kg in 100 µl PBS solution by i.v. injections in the tail vein. Animals were monitored regularly for changes in body weight and body conditions. The study was terminated on day 48 post-treatment when animals began to exhibit severe hunched posture and signs of gait rigidity. Tumor tissues were collected, and mass and weight were recorded *ex vivo*. The formula, volume = 1/2 (length × width^2^), was used to calculate the tumor volumes.

#### Biodistribution

SKOV-3 tumor-bearing BALB/c nude mice were administered a single intravenous dose of 10 mg/kg of the AFC (100 µL, PBS solution) on day 60 following tumor cell inoculation. Animals were sacrificed at predetermined time points—1 hour, 1 day, 3 days, 5 days, and 7 days post-injection (n=3 per time point). Major organs were harvested, and fluorescence signals were quantified *ex vivo* using an IVIS to assess the biodistribution of the AFC. Untreated animal was used as a control.

## Availability of data and materials

The raw data required to reproduce these findings are available from the authors upon request.

## Author Contributions

Conceptualization: AR, MWG Methodology: AR, TYS Investigation: AR, TYS Visualization: AR Supervision: MWG Writing—original draft: AR Writing—review & editing: All authors Funding: MWG, AR, TYS

## Acknowledgements

We acknowledge and thank the National Institutes of Health, PhRMA Foundation, Chemical Instrumentation Center at Boston University, and Boston University

## REFERENCES

1. Winssinger N. Bioorthogonal chemistry. Chimia (Aarau). 2018;72(11):A755.

2. Agard NJ, Prescher JA, Bertozzi CR. A strain-promoted [3 + 2] azide-alkyne cycloaddition for covalent modification of biomolecules in living systems. J Am Chem Soc. 2004;126(46):15046–7.

3. You J, Zhang J, Wang J, Jin M. Cysteine-Based Coupling: Challenges and Solutions. Bioconjug Chem. 2021;32(8):1525–34.

4. Fu Z, Li S, Han S, Shi C, Zhang Y. Antibody drug conjugate: the “biological missile” for targeted cancer therapy. Vol. 7, Signal Transduction and Targeted Therapy. Springer Nature; 2022.

5. Gordon MR, Canakci M, Li L, Zhuang J, Osborne B, Thayumanavan S. Field Guide to Challenges and Opportunities in Antibody-Drug Conjugates for Chemists. Bioconjug Chem. 2015;26(11):2198–215.

6. Lazar AC, Wang L, Blättler WA, Amphlett G, Lambert JM, Zhang W. Analysis of the composition of immunoconjugates using size-exclusion chromatography coupled to mass spectrometry. Rapid Commun Mass Spectrom. 2005;19(13):1806–14.

7. Wu G, Gao Y, Liu D, Tan X, Hu L, Qiu Z, et al. Study on the heterogeneity of T-DM1 and the analysis of the unconjugated linker structure under a stable conjugation process. ACS Omega. 2019;4(5):8834–45.

8. Spears RJ, Chudasama V. Recent advances in N- and C-terminus cysteine protein bioconjugation. Curr Opin Chem Biol [Internet]. 2023;75:102306. Available from: 10.1016/j.cbpa.2023.102306

9. Tantipanjaporn A, Wong MK. Development and Recent Advances in Lysine and N- Terminal Bioconjugation for Peptides and Proteins. Molecules. 2023;28(3).

10. Wang Y, Zhang J, Han B, Tan L, Cai W, Li Y, et al. Noncanonical amino acids as doubly bio-orthogonal handles for one-pot preparation of protein multiconjugates. Nat Commun. 2023;14(1).

11. Kim CH, Axup JY, Schultz PG. Protein conjugation with genetically encoded unnatural amino acids. Curr Opin Chem Biol. 2013;17(3):412–9.

12. Maza JC, McKenna JR, Raliski BK, Freedman MT, Young DD. Synthesis and incorporation of unnatural amino acids to probe and optimize protein bioconjugations. Bioconjug Chem. 2015;26(9):1884–9.

13. Jang S, Sachin K, Lee HJ, Kim DW, Lee HS. Development of a simple method for protein conjugation by copper-free click reaction and its application to antibody-free western blot analysis. Bioconjug Chem. 2012;23(11):2256–61.

14. Jaramillo ML, Sulea T, Durocher Y, Acchione M, Schur MJ, Robotham A, et al. A glyco-engineering approach for site-specific conjugation to Fab glycans. MAbs [Internet]. 2023;15(1):1–15. Available from: 10.1080/19420862.2022.2149057

15. Zhou Q, Stefano JE, Manning C, Kyazike J, Chen B, Gianolio DA, et al. Site-Specific Antibody − Drug Conjugation through Glycoengineering. 2014;

16. Li X, Fang T, Boons G. Preparation of Well-Defined Antibody–Drug Conjugates through Glycan Remodeling and Strain-Promoted Azide–Alkyne Cycloadditions. Angew Chemie. 2014;126(28):7307–10.

17. Badescu G, Bryant P, Bird M, Henseleit K, Swierkosz J, Parekh V, et al. Bridging disulfides for stable and defined antibody drug conjugates. Bioconjug Chem. 2014;25(6):1124–36.

18. Kuan SL, Wang T, Weil T. Site-Selective Disulfide Modification of Proteins: Expanding Diversity beyond the Proteome. Chem - A Eur J. 2016;22(48):17112–29.

19. Gautier A, Hinner MJ. Site-Specific Protein Labeling: Methods and Protocols. Site-Specific Protein Labeling Methods Protoc. 2015;1266:1–267.

20. Popp MWL, Ploegh HL. Making and breaking peptide bonds: Protein engineering using sortase. Angew Chemie - Int Ed. 2011;50(22):5024–32.

21. Debon A, Siirola E, Snajdrova R. Enzymatic Bioconjugation: A Perspective from the Pharmaceutical Industry. JACS Au. 2023;3(5):1267–83.

22. Milczek EM. Commercial Applications for Enzyme-Mediated Protein Conjugation: New Developments in Enzymatic Processes to Deliver Functionalized Proteins on the Commercial Scale. Chem Rev. 2018;118(1):119–41.

23. Pirzer T, Becher KS, Rieker M, Meckel T, Mootz HD, Kolmar H. Generation of Potent Anti-HER1/2 Immunotoxins by Protein Ligation Using Split Inteins. ACS Chem Biol. 2018;13(8):2058–66.

24. Lieser RM, Yur D, Sullivan MO, Chen W. Site-Specific Bioconjugation Approaches for Enhanced Delivery of Protein Therapeutics and Protein Drug Carriers. Bioconjug Chem. 2020;31(10):2272–82.

25. Reddington SC, Howarth M. Secrets of a covalent interaction for biomaterials and biotechnology: SpyTag and SpyCatcher. Curr Opin Chem Biol [Internet]. 2015;29:94–9. Available from: 10.1016/j.cbpa.2015.10.002

26. Lotze J, Reinhardt U, Seitz O, Beck-Sickinger AG. Peptide-tags for site-specific protein labelling: In vitro and in vivo. Mol Biosyst. 2016;12(6):1731–45.

27. Wang R, Hu B, Pan Z, Mo C, Zhao X, Liu G, et al. Antibody–Drug Conjugates (ADCs): current and future biopharmaceuticals. J Hematol Oncol [Internet]. 2025;18(1). Available from: 10.1186/s13045-025-01704-3

28. Jian A, Zhao G, Zhou J, Wang S, Li N. How to design next-generation of antibody-drug conjugates for cancer treatment: Lessons from unsuccessful clinical trials. Cancer Lett [Internet]. 2025;623(November 2024):217535. Available from: 10.1016/j.canlet.2025.217535

29. Webber MJ, Langer R. Drug delivery by supramolecular design. Chem Soc Rev. 2017;46(21):6600–20.

30. Bernhard S, Tibbitt MW. Supramolecular engineering of hydrogels for drug delivery. Adv Drug Deliv Rev. 2021;171:240–56.

31. Mattia E, Otto S. Supramolecular systems chemistry. Nat Nanotechnol. 2015;10(2):111–9.

32. Stupp SI, Palmer LC. Supramolecular chemistry and self-assembly in organic materials design. Chem Mater. 2014;26(1):507–18.

33. Webber MJ, Appel EA, Meijer EW, Langer R. Supramolecular biomaterials. Nat Mater. 2015;15(1):13–26.

34. Thomas F, Boyle AL, Burton AJ, Woolfson DN. A set of de novo designed parallel heterodimeric coiled coils with quantified dissociation constants in the micromolar to sub-nanomolar regime. J Am Chem Soc. 2013;135(13):5161–6.

35. Utterström J, Naeimipour S, Selegård R, Aili D. Coiled coil-based therapeutics and drug delivery systems. Adv Drug Deliv Rev. 2021;170:26–43.

36. Aschmann D, Knol RA, Kros A. Lipid-Based Nanoparticle Functionalization with Coiled-Coil Peptides for In Vitro and In Vivo Drug Delivery. Acc Chem Res. 2024;57(8):1098–110.

37. Britton D, Sun JW, Renfrew PD, Montclare JK. Design of Coiled-Coil Protein Nanostructures for Therapeutics and Drug Delivery. Annu Rev Chem Biomol Eng. 2024;15(1):25–50.

38. Wu Y, Collier JH. α-Helical coiled-coil peptide materials for biomedical applications. Wiley Interdiscip Rev Nanomedicine Nanobiotechnology. 2017;9(2):1–17.

39. Britton D, Legocki J, Paul D, Katsara O, Aristizabal O, Pandya N, et al. Coiled-Coil Protein Hydrogels Engineered with Minimized Fiber Diameters for Sustained Release of Doxorubicin in Triple-Negative Breast Cancer. ACS Biomater Sci Eng. 2024;10(5):3425–37.

40. Bennett JI, Boit MOK, Gregorio NE, Zhang F, Kibler RD, Hoye JW, et al. Genetically Encoded XTEN-based Hydrogels with Tunable Viscoelasticity and Biodegradability for Injectable Cell Therapies. Adv Sci. 2024;11(24):1–11.

41. Gil-Garcia M, Ventura S. Multifunctional antibody-conjugated coiled-coil protein nanoparticles for selective cell targeting. Acta Biomater. 2021;131:472–82.

42. Zeng Y, Estapé Senti M, Labonia MCI, Papadopoulou P, Brans MAD, Dokter I, et al. Fusogenic Coiled-Coil Peptides Enhance Lipid Nanoparticle-Mediated mRNA Delivery upon Intramyocardial Administration. ACS Nano. 2023;17(23):23466–77.

43. Lainšček D, Forstnerič V, Mikolič V, Malenšek Š, Pečan P, Benčina M, et al. Coiled-coil heterodimer-based recruitment of an exonuclease to CRISPR/Cas for enhanced gene editing. Nat Commun. 2022;13(1).

44. Wang J, Yu Y, Xia J. Short peptide tag for covalent protein labeling based on coiled coils. Bioconjug Chem. 2014;25(1):178–87.

45. Liu D, Zhao J, Song Y. Engineering switchable and programmable universal CARs for CAR T therapy. J Hematol Oncol. 2019;12(1):1–9.

46. Bell M, Lange S, Sejdiu BI, Ibanez J, Shi H, Sun X, et al. Modular chimeric cytokine receptors with leucine zippers enhance the antitumour activity of CAR T cells via JAK/STAT signalling. Nat Biomed Eng. 2024;8(4):380–96.

47. Gradišar H, Jerala R. De novo design of orthogonal peptide pairs forming parallel coiled-coil heterodimers. J Pept Sci. 2011;17(2):100–6.

48. Coin I, Beyermann M, Bienert M. Solid-phase peptide synthesis: From standard procedures to the synthesis of difficult sequences. Nat Protoc. 2007;2(12):3247–56.

49. Isidro-Llobet A, Kenworthy MN, Mukherjee S, Kopach ME, Wegner K, Gallou F, et al. Sustainability Challenges in Peptide Synthesis and Purification: From R&D to Production. J Org Chem. 2019 Apr 19;84(8):4615–28.

50. Zimmerman ES, Heibeck TH, Gill A, Li X, Murray CJ, Madlansacay MR, et al. Production of site-specific antibody-drug conjugates using optimized non-natural amino acids in a cell-free expression system. Bioconjug Chem. 2014;25(2):351–61.

51. Adhikari A, Bhattarai BR, Aryal A, Thapa N, Kc P, Adhikari A, et al. Reprogramming natural proteins using unnatural amino acids. RSC Adv. 2021;11(60):38126–45.

52. Ahn JH, Kang S, Park S, Song H, Yun Y, Choi S, et al. Reversible Protein Conjugation on Live Cell Surfaces by Specific Recognition between Coiled-Coil Motifs of Natural Amino Acid Sequences. Biomacromolecules. 2020;21(9):3539–46.

53. Bose S, Sharma S, Kumar A, Mishra Y, Mishra V. Ovarian cancer and its management through advanced drug delivery system. Med Oncol [Internet]. 2025;42(3):1–16. Available from: 10.1007/s12032-025-02621-8

54. Wang Z, Meng F, Zhong Z. Emerging targeted drug delivery strategies toward ovarian cancer. Adv Drug Deliv Rev [Internet]. 2021;178:113969. Available from: 10.1016/j.addr.2021.113969

55. Nogueira-Rodrigues A, Giannecchini GV, Secord AA. Real world challenges and disparities in the systemic treatment of ovarian cancer. Gynecol Oncol [Internet]. 2024;185:180–5. Available from: 10.1016/j.ygyno.2024.02.021

56. Konstantinopoulos PA, Matulonis UA. Clinical and translational advances in ovarian cancer therapy. Nat Cancer. 2023;4(9):1239–57.

57. Dubowchik GM, Firestone RA, Padilla L, Willner D, Hofstead SJ, Mosure K, et al. Cathepsin B-labile dipeptide linkers for lysosomal release of doxorubicin from internalizing immunoconjugates: Model studies of enzymatic drug release and antigen-specific in vitro anticancer activity. Bioconjug Chem. 2002;13(4):855–69.

58. Wakankar A, Chen Y, Gokarn Y, Jacobson FS. Analytical methods for physicochemical characterization of antibody drug conjugates. MAbs. 2011;3(2):161–72.

59. Hamblett KJ, Senter PD, Chace DF, Sun MMC, Lenox J, Cerveny CG, et al. Effects of Drug Loading on the Antitumor Activity of a Monoclonal Antibody Drug Conjugate. Clin Cancer Res. 2004;10(425):7063–70.

60. Kuo WY, Lin JJ, Hsu HJ, Chen H Sen, Yang AS, Wu CY. Noninvasive assessment of characteristics of novel anti-HER2 antibodies by molecular imaging in a human gastric cancer xenograft-bearing mouse model. Sci Rep [Internet]. 2018;8(1):1–9. Available from: 10.1038/s41598-018-32094-x

61. Park S, Nedrow JR, Josefsson A, Sgouros G. Human HER2 overexpressing mouse breast cancer cell lines derived from MMTV.f.HuHER2 mice: characterization and use in a model of metastatic breast cancer. Oncotarget. 2017;8(40):68071–82.

62. Galsky MD, Grande E, Necchi A, Koontz MZ, Iyer G, Campbell MT, et al. Phase 3 open-label, randomized, controlled study of disitamab vedotin with pembrolizumab versus chemotherapy in patients with previously untreated locally advanced or metastatic urothelial carcinoma that expresses HER2 (DV-001). J Clin Oncol. 2024;42(4_suppl):TPS717–TPS717.

63. Shi F, Liu Y, Zhou X, Shen P, Xue R, Zhang M. Disitamab vedotin: a novel antibody-drug conjugates for cancer therapy. Drug Deliv [Internet]. 2022;29(1):1335–44. Available from: 10.1080/10717544.2022.2069883

64. Choy N, Raussens V, Narayanaswami V. Inter-molecular coiled-coil formation in human apolipoprotein E C-terminal domain. J Mol Biol. 2003;334(3):527–39.

