## Supporting Information for "A Supramolecular Self-assembly Approach to Site-Specific Antibody Conjugates *via* a Coiled-coil Peptides Platform"

**Table SI1.** Sequences of orthogonal peptides which form coiled-coils structures P1:P2, P3:P4, and $P1N_3$:$P2N_3$, $P3N_3$:$P4N_3$, $P3_{5mer}$:$P4_{5mer}N_3$, $P3_{5mer}$-cys:$P4_{5mer}$-cys$N_3$.

| | Sequence | Use | Sequence source |
|---|---|---|---|
| **P1** | SPED EIQALEE ENAQLEQ ENAALEE EIAQLEY G | receiving peptide | (1) |
| **P2** | SPED KIAQLKE KNAALKE KNQQLKE KIQALKY G | receiving peptide | (1) |
| **P3** | SPED EIQQLEE EIAQLEQ KNAALKE KNQALKY G | receiving peptide | (1) |
| **P4** | SPED KIAQLKQ KIQALKQ ENQQLEE ENAALEY G | receiving peptide | (1) |
| **$P1N_3$** | SPED EIQALEE ENAQLEQ ENAALEE EIAQLEY G(K-azido)G | docking peptide | This paper, modified from (1) |
| **$P2N_3$** | SPED KIAQLKE KNAALKE KNQQLKE KIQALKY G(K-azido)G | docking peptide | This paper, modified from (1) |
| **$P3N_3$** | SPED EIQQLEE EIAQLEQ KNAALKE KNQALKY G(K-azido)G | docking peptide | This paper, modified from (1) |
| **$P4N_3$** | SPED KIAQLKQ KIQALKQ ENQQLEE ENAALEY G(K-azido)G | docking peptide | This paper, modified from (1) |
| **$P3_{5mer}$** | SPED EIQQLEE EIAQLEQ KNAALKE KNQALKY EIQQLEE | receiving peptide | This paper, modified from (1) |
| **$P3_{5mer}$-cys** | SPED EIQQLEE EIAQLEQ KNAALKE KNQALKY E**C**QQLEE G | receiving peptide | This paper, modified from (1) |
| **$P4_{5mer}N_3$** | SPED KIAQLKQ KIQALKQ ENQQLEE ENAALEY KIAQLKQ G(K-azido)G | docking peptide | This paper, modified from (1) |
| **$P4_{5mer}$-cys$N_3$** | SPED KIAQLKQ KIQALKQ ENQQLEE ENAALEY K**C**AQLKQ G(K-azido)G | docking peptide | This paper, modified from (1) |

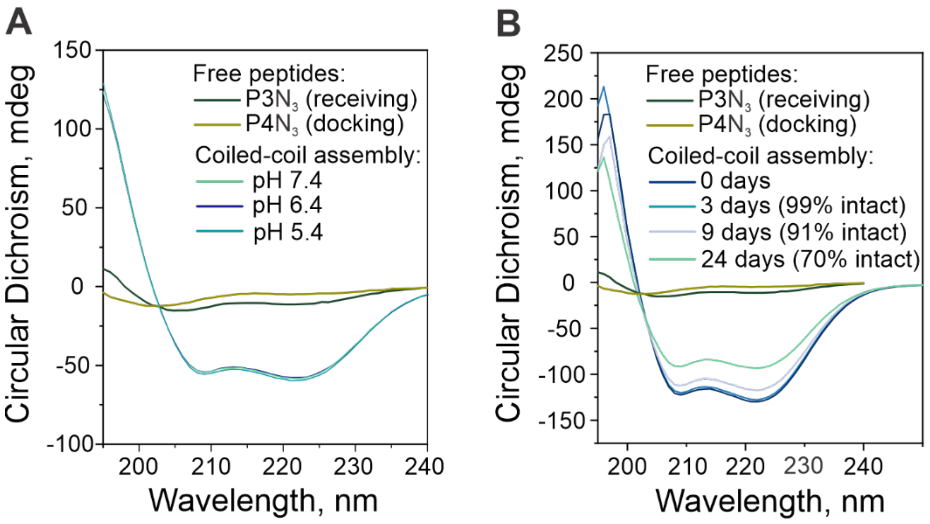


**Figure SI1.** Circular Dichroism (CD) spectra of individual orthogonal P3 and P4 peptides modified with azide ($N_3$) functional group and their equimolar mixture **A.** at different pH demonstrating a robust coiled-coil formation and **B.** over time after 37 °C incubation. The proportion of intact coiled-coil assemblies relative to single α-helices was estimated by calculating the ratio of ellipticity at 222 nm to that at 208 nm (2).

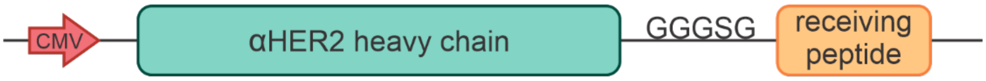


**Figure SI2.** Plasmid scheme of the heavy chain of the trastuzumab antibody fused to the receiving peptide at the C-terminal end, cloned into the pcDNA3.1(-) vector.

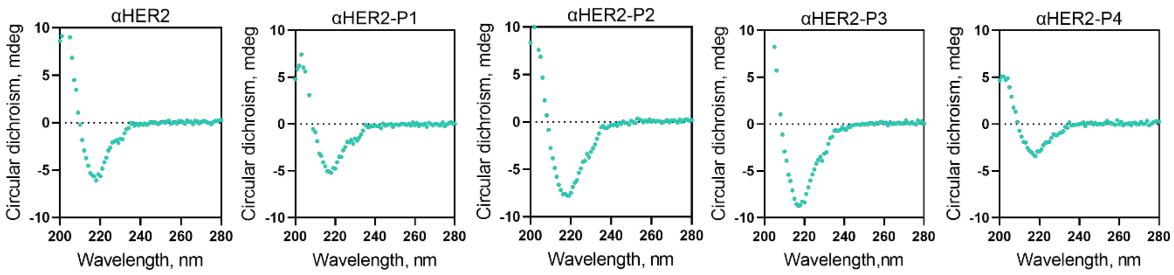


**Figure SI3.** Circular dichroism (CD) spectra of native mAb (αHER2, trastuzumab) and antibodies fused to receiving peptides P1, P2, P3 or P4 measured in PBS buffer pH=7.4 at room temperature.

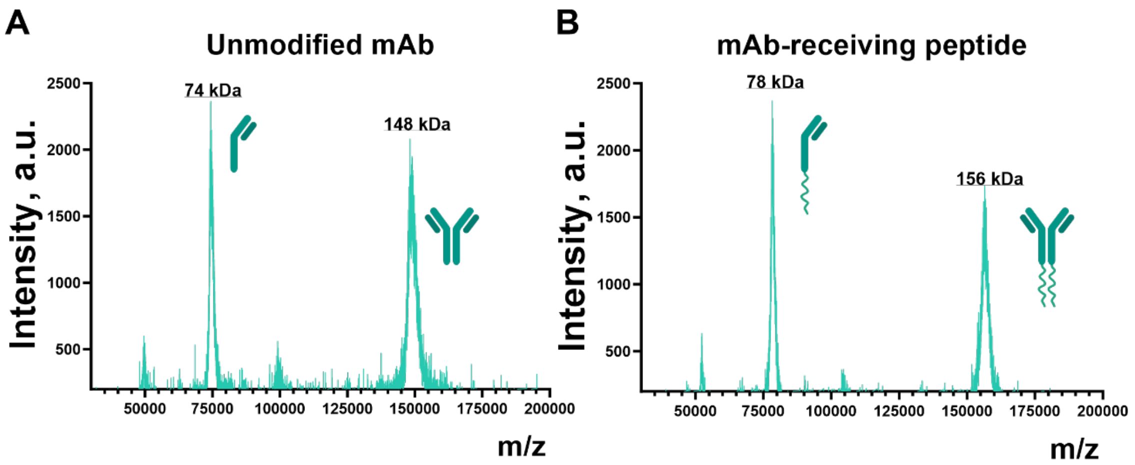


**Figure SI4.** MALDI-TOF spectra of **A.** native trastuzumab mAb (αHER2) and **B.** trastuzumab antibodies fused to receiving peptides P3 (αHER2-P3), where the increase in molecular weight corresponds to the mass of two P3 peptides (P3 $M_w$= 4 kDa each) fused to the heavy chain of the antibody.

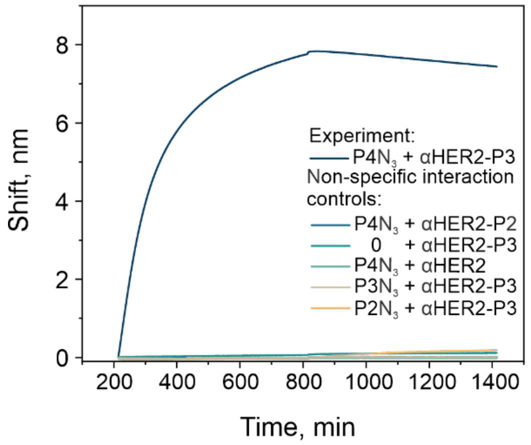


**Figure SI5.** Binding affinity of αHER2-P3 to $P4N_3$ peptide biotinylated with DBCO-PEG$_4$-biotin measured by biolayer interferometry (experimental curve) and specificity controls: 1) target docking peptide and mAb fused to non-specific receiving peptide ($P4N_3$ + αHER2-P2); 2) no docking peptide and mAb fused to specific receiving peptide (0 + αHER2-P3); 3) target docking peptide and native mAb ($P4N_3$ + αHER2); 4) self-interaction control - receiving peptide and mAb fused to the same receiving peptide ($P3N_3$ + αHER2-P3); 5) non-specific docking peptide and mAb fused to target receiving peptide ($P2N_3$ + αHER2-P3).

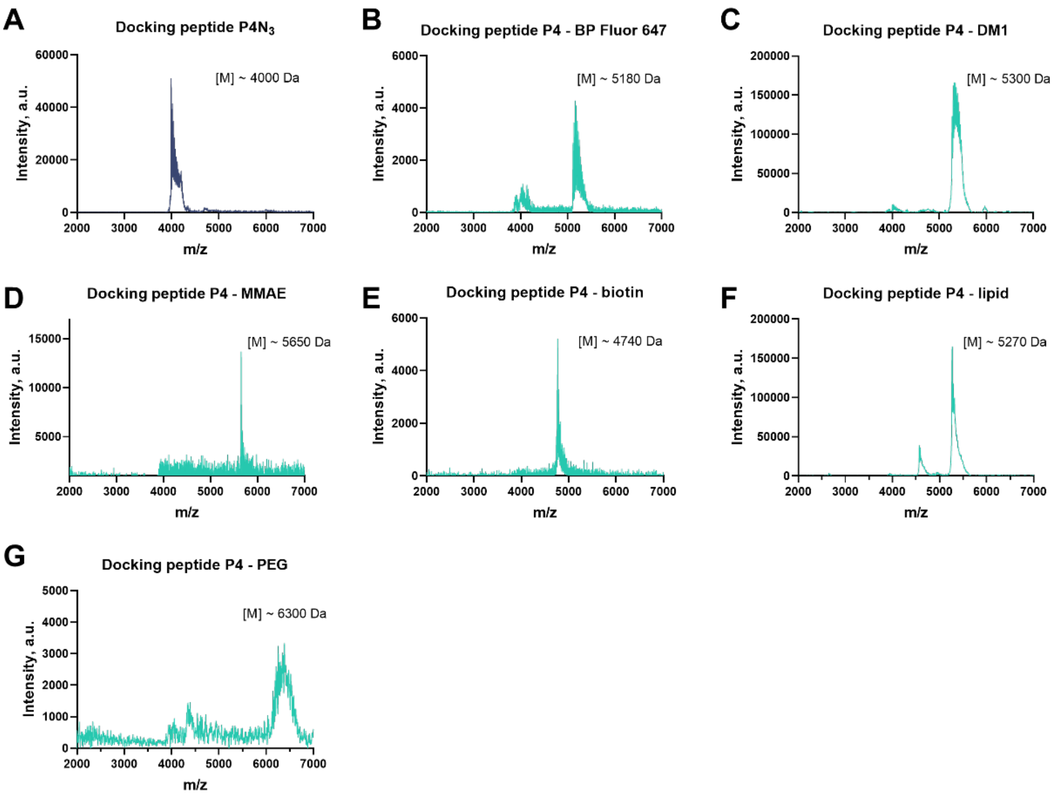


**Figure SI6.** MALDI-TOF spectra of **A.** unmodified docking peptide $P4N_3$ and $P4N_3$ docking peptide conjugated to **B.** BP Fluor 647 DBCO ($M_w$ = 1117.3 g/mol), **C.** DBCO-$PEG_4$-DM1 ($M_w$ = 1272.9 g/mol), **D.** DBCO-$PEG_4$-Val-Cit-PAB-MMAE ($M_w$ = 1658.1 g/mol), **E.** DBCO-$PEG_4$-biotin ($M_w$ = 749.92 g/mol), **F.** DBCO-$PEG_4$-DSPE ($M_w$ = 1282.7 g/mol), **G.** DBCO-$PEG_{2k}$-OH ($M_w$ ≈ 2322 g/mol).

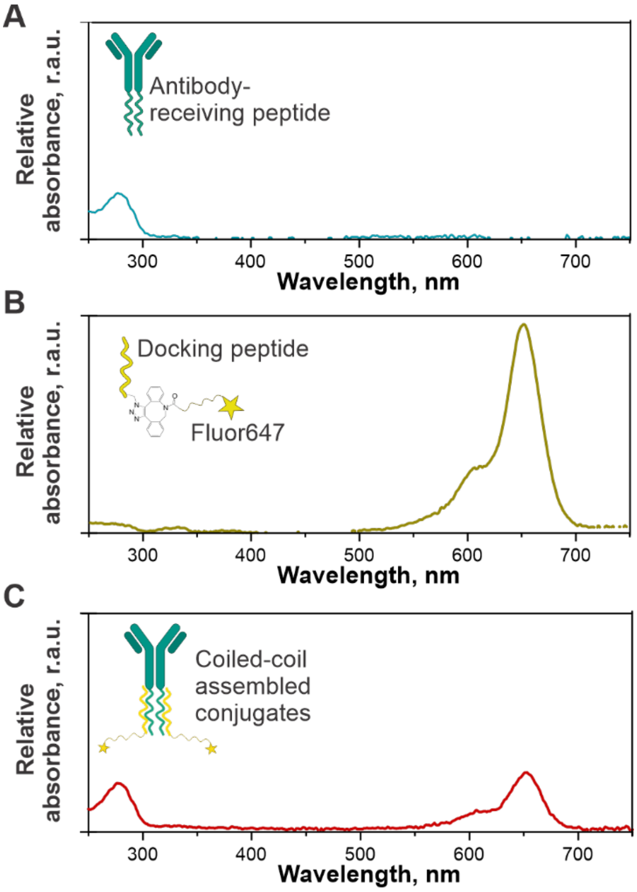


**Figure SI7.** UV-Vis analyses of **A.** αHER2-P3, **B.** DBCO-modified BP Fluor 647-loaded docking peptide P4 and **C.** coiled-coil mediated αHER2-Fluor647 conjugate. Fluorophore-antibody ratio (FAR) of 2 was calculated using the following equation and standard curves to determine extinction coefficients for each assembling part:

$$FAR = \frac{\varepsilon_{Ab}^{647} - R\varepsilon_{Ab}^{280}}{R\varepsilon_{D}^{280} - \varepsilon_{D}^{647}}, \text{ where } R = {}^{A_{647}}/_{A_{280}}$$

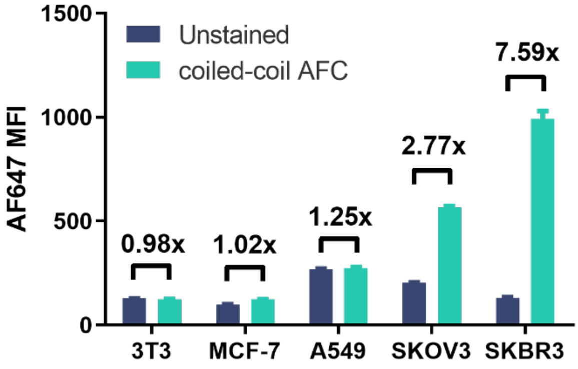


**Figure SI8.** Flow cytometry analysis of the mean fluorescence intensity of cells expressing different level of ErbB2/Her2 using coiled-coil formed αHER2-Fluor647 antibodies. Coiled-coil based AFC target ErbB2/Her2 positive SKOV-3 and SKBR-3 cells while do not show significant targeting of ErbB2/Her2 low or negative cells (NIH 3T3, MCF-7 and A549).

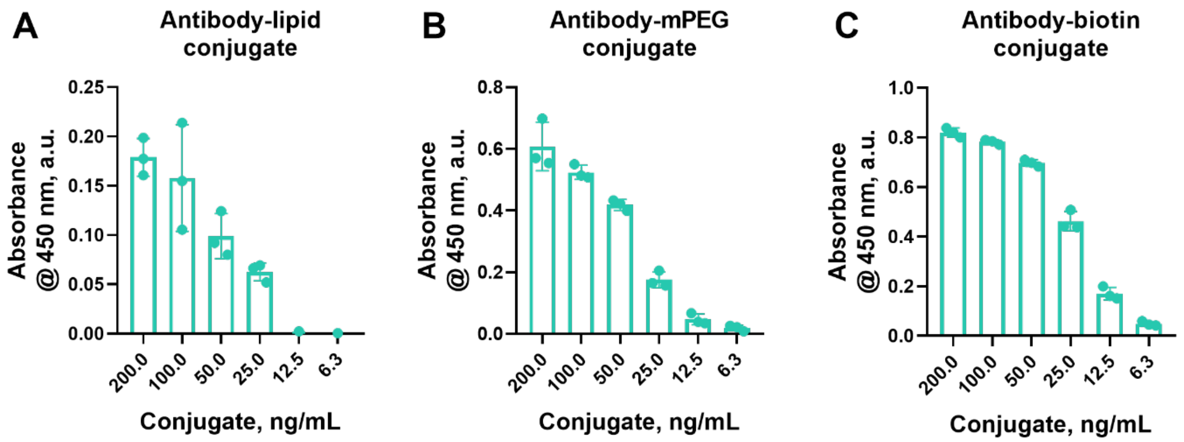


**Figure SI9.** Indirect ELISA for detection of payloads on antibody conjugates using ErbB2/Her2 antigen and payload-specific secondary antibodies. **A.** Antibody-lipid conjugates (DSPE-PEG$_{5k}$-DBCO) detected with anti-PEG HRP secondary antibodies **B.** Antibody-mPEG conjugates (DBCO-mPEG$_{10k}$) detected with anti-PEG HRP secondary antibodies and **C.** Antibody-biotin conjugates (DBCO-PEG$_4$-biotin) detected with streptavidin HRP.

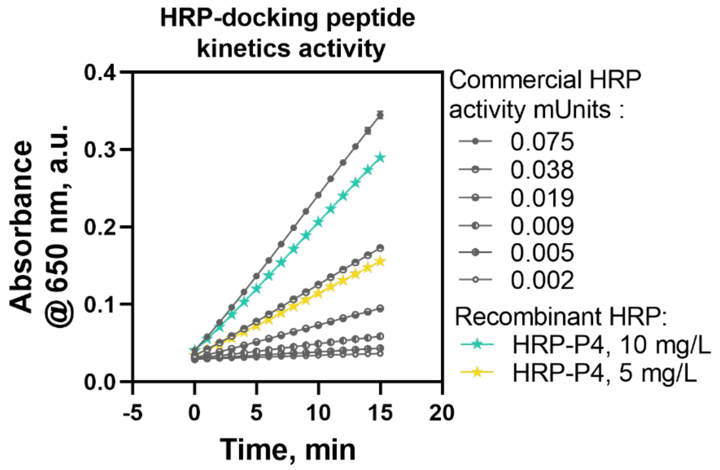


**Figure SI10.** Kinetic TMB-based HRP activity assay measuring the activity of recombinant HRP-P4 conjugate compared to serial dilution of commercially available native HRP (Sigma, CAS #9003-99-0). Absorbance measured at 650 nm with 1 minute interval.

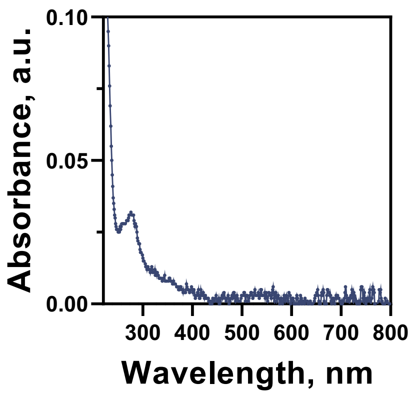


**Figure SI11.** UV-Vis analysis of antibody αHER2-P3 reacted with a fluorophore labeled polyT$_{15}$CCC-TAMRA oligo demonstrating the absence of TAMRA fluorophore in the final product post purification.

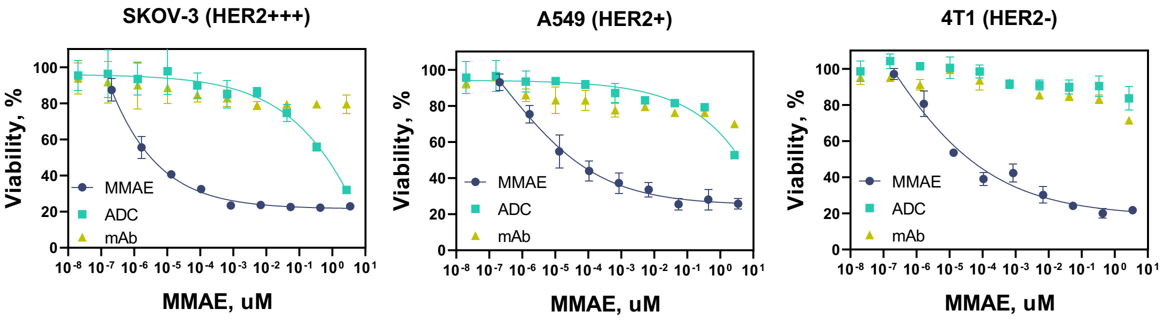


**Figure SI12.** Cytotoxicity MTS assay of the coiled-coil peptides created αHER2 ADC bearing MMAE drug against: **A.** SKOV-3 –high HER2 expression cell line; **B.** A549 – low HER2 expression and **C.** 4T1 – HER2 negative cells; compared to equivalent dose of free MMAE drug and unloaded αHER2-P3 antibody over 72 hours.

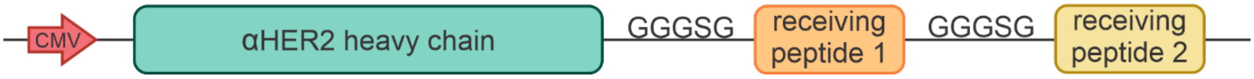


**Figure SI13.** Plasmid scheme of the trastuzumab antibody heavy chain fused to two sequential orthogonal coil receiving peptides, cloned in the pcDNA3.1(-) vector.

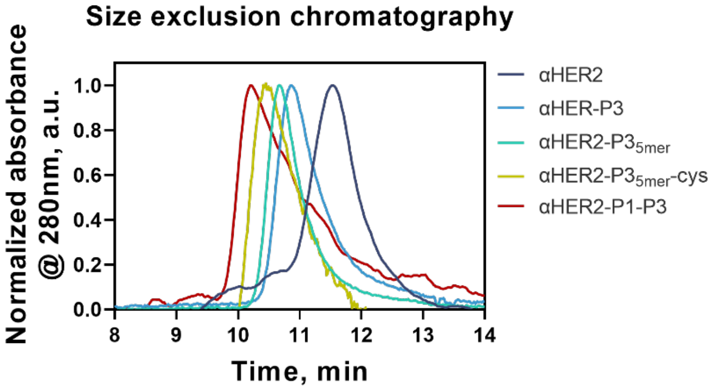


**Figure SI14.** SEC-UV traces at 280 nm of native trastuzumab (αHER2), trastuzumab fused to different receiving peptides (αHER2-P3, αHER2-P3$_{5mer}$, and αHER2-P3$_{5mer}$-cys) and trastuzumab fused to two sequential orthogonal receiving peptides (αHER2-P1-P3) showing the shortening retention time depending on the molecular weight of the sample.

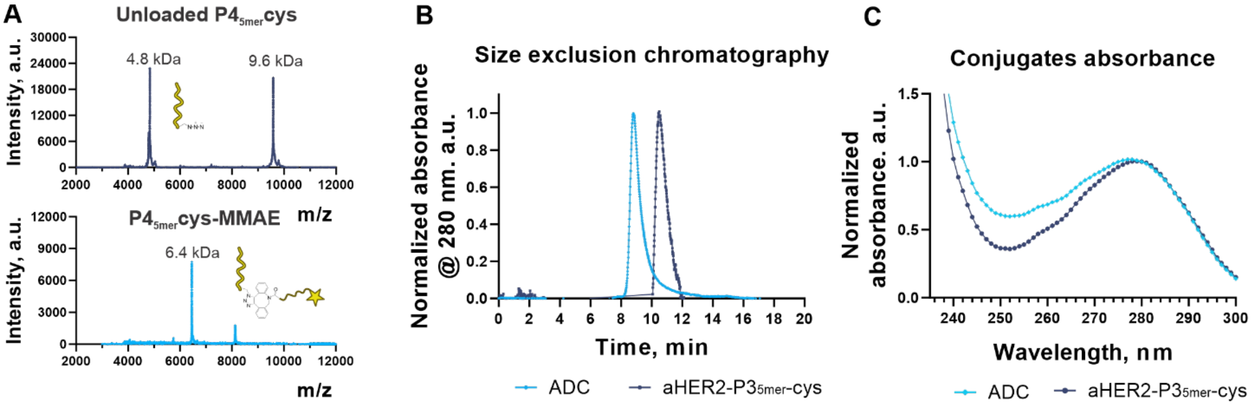


**Figure SI15. ADC preparation. A.** MALDI-TOF spectra of unmodified docking peptide P4$_{5mer}$-cys-$N_3$ ($M_w$=4.8 kDa) and docking peptide conjugated to cleavable-linker-MMAE P4$_{5mer}$-cys-MMAE ($M_w$=6.4 kDa), where the increase in molecular weight corresponds to the mass of one DBCO-Val-Cit-PAB-MMAE molecule ($M_w$ = 1.6 kDa), **B.** SEC-UV traces at 280 nm of unloaded mAb (αHER2-P3$_{5mer}$-cys) and fully assembled ADC showing the shortening retention based on increasing the molecular weight due to conjugation to P4$_{5mer}$-cys-MMAE, **C.** An overlay of UV-vis spectra of the starting mAb and conjugated ADC, normalized to the 280 nm absorbance, showing the increase in the 248 nm absorbance due to drug conjugation as described in (3).

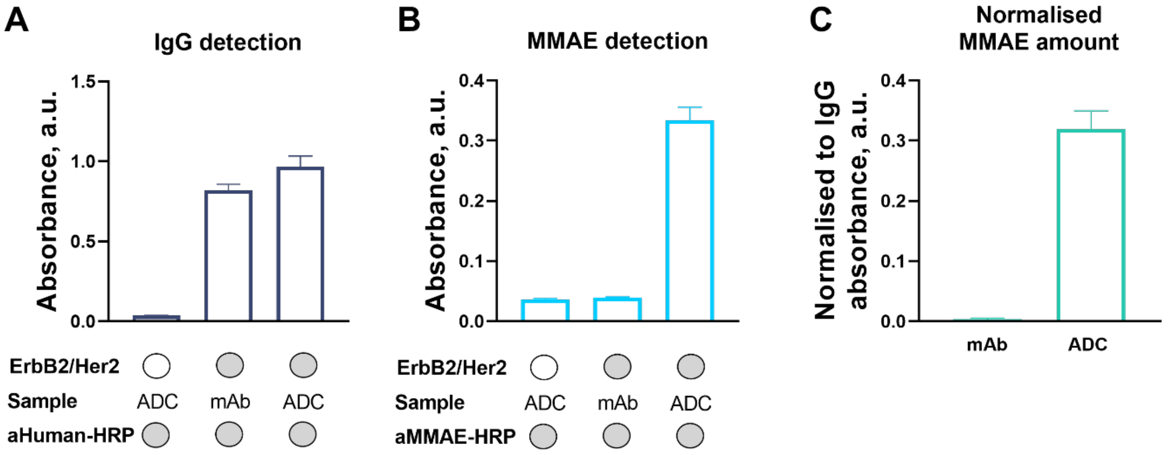


**Figure SI16.** Indirect ELISA-based assessment of MMAE conjugation to aHER2-P3$_{5mer}$-cys, **A.** Absorbance signal at 450 nm obtained using anti-human secondary detection antibodies, **B.** Absorbance signal at 450 nm obtained using anti-MMAE secondary detection antibodies to detect conjugated MMAE, **C.** Normalized MMAE detection signal relative to the detected antibody level, indicating MMAE conjugation in the final ADC.

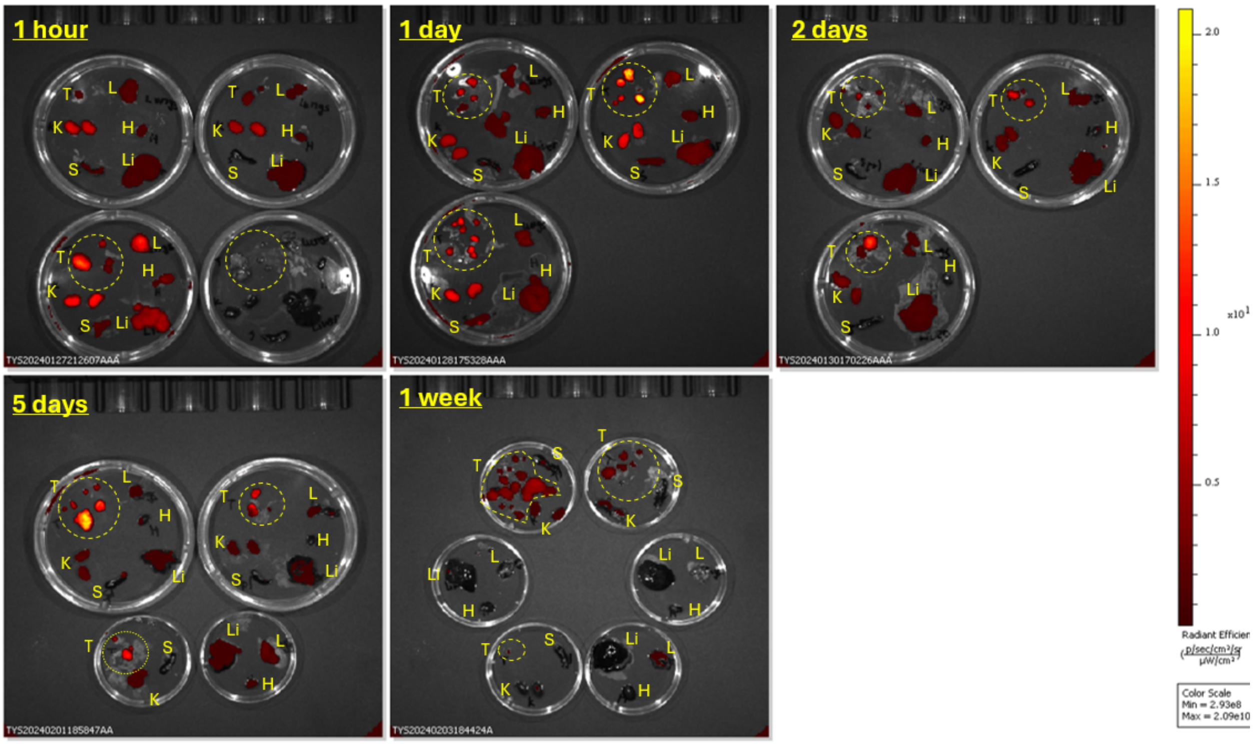


**Figure SI17**. *Ex vivo* organ biodistribution of coiled-coil AFC injected i.v. at 10 mg/kg dose into SKOV-3 tumor bearing NSG mice at different time points (n=3 per time point). The total radiant efficiency was calculated using IVIS software keeping area of ROI the same across all the samples. Letters indicate: T – tumor (dashed outline), L – lungs, H – heart, Li – liver, S – spleen, K – kidney

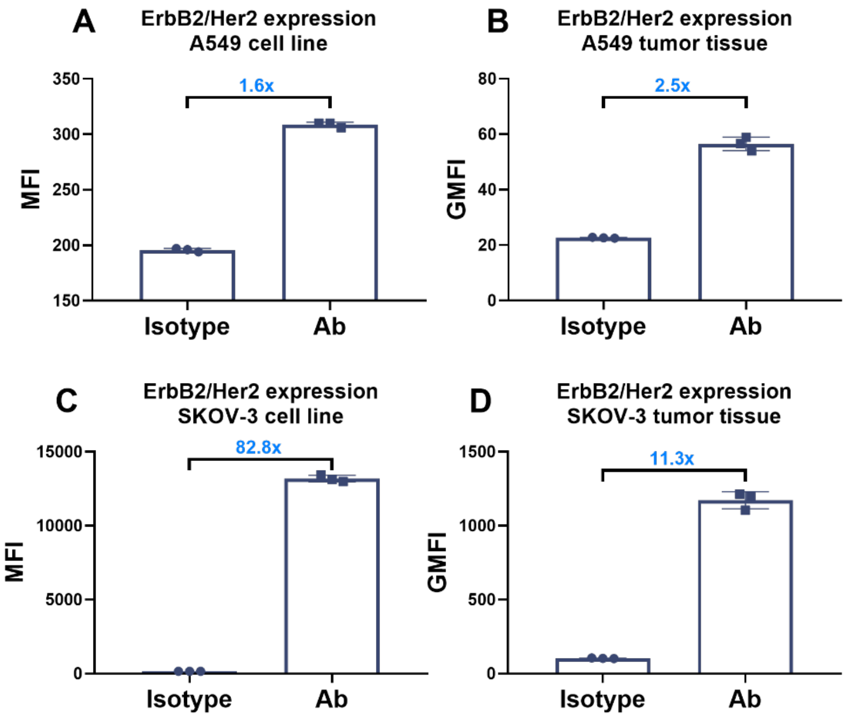


**Figure SI18.** Flow cytometry evaluation of αHER2-Fluor647 binding to ErbB2/Her2 on A549 and SKOV-3 at cell line and tumor tissue levels.

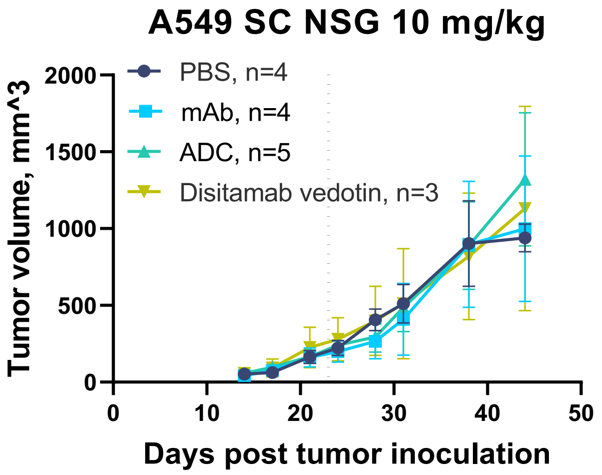


**Figure SI19.** Average tumors size after different treatments in female NSG mice engrafted subcutaneously with 2 million A549 tumor cells and treated on day 23 (average tumor volume ~175 mm$^3$) with either PBS or coiled-coil ADC or mAb (αHER2-P3$_{5mer}$-cys) or commercial ADC (disitamab vedotin) at the dose 10 mg/kg. The dashed line indicates the treatment day.

### SI METHODS

**Efficacy study in ErbB2/Her2 low animal model.** Female NSG mice (6–8 weeks old) were obtained from the Jackson Laboratory (#005557). The xenograft model of A549 was established by injecting A549 cells (2 × 10$^6$ cells in 0.2 ml PBS) subcutaneously. On day 23 post-inoculation, when the average tumor volume reached ~175 mm$^3$, mice were randomized into treatment groups. Mice were administered a single dose of 100 µl of PBS or coiled-coil ADC, disitamab vedotin or unloaded mAb (αHER2-P3$_{5mer}$-cys) at a dose of 10 mg/kg in 100 µl of PBS solution by i.v. injections in the tail vein. Animals were monitored regularly for changes in body weight, body conditions and tumor volume. The formula, volume = 1/2 (length × width$^2$), was used to calculate the tumor volumes.

**Polymer, lipid, biotin and MMAE conjugation assessment assay.** Activity of the αHER2-mPEG, αHER2-DSPE lipid or αHER2-biotin conjugates was assessed by indirect ELISA. Nunc MaxiSorp 96-well ELISA plates were coated overnight at 4 °C with 2 µg/mL of recombinant human ErbB2/Her2 protein (Cat. #10126-ER, R&D Systems Inc.) in 0.05 M carbonate–bicarbonate buffer (pH 9.6). Plates were then blocked with 5% bovine serum albumin (BSA) in PBS and washed with PBS containing 0.05% Tween-20 (PBS-T) between each step, following a standard ELISA protocol. The αHER2-payload conjugates were serially diluted 2-fold in blocking buffer (starting from 200 ng/mL diluting to 6.3 ng/mL) and incubated in ELISA plate for 1 hour at 37 °C. After washing steps, the payload was detecting using anti-payload antibodies (500 ng/mL anti-polyethylene glycol [PEG-B-47] HRP, abcam) or streptavidin-HRP (1:40 dilution, R&D Systems Inc.). 60 µL TMB solution (1-Step™ TMB ELISA Substrate Solutions, Thermo Scientific™) was used as HRP substrate for detection. The reaction was stopped with 60 µL of 2N sulfuric acid and the absorbance at 450 nm was measured using a microplate reader (SpectraMax iD3, Molecular devices). Wells with no plated ErbB2/Her2 or no conjugates were used as controls. The MMAE conjugation to the aHER2-P3$_{5mer}$-cys for the efficacy study was assessed with similar indirect ELISA. Unloaded aHER2-P3$_{5mer}$-cys were used as an additional control and anti-humman IgG (1:5000 dilution Goat anti-human IgG H&L HRP, abcam) or anti-MMAE secondary antibodies (100 ng/mL HRP conjugated Monoclonal Anti-MMAE&MMAF Antibody, ACROBiosystems) were used for detection.

**Flow cytometry.** The assessment of αHER2-Fluor647 binding to the cells ErbB2/Her2 as well as ErbB2/Her2 expression was performed by flow cytometry using the Attune NxT Flow Cytometer (Thermo Fisher Scientific) and analyzed using FlowJo (v10.8.1). For αHER2-Fluor647 binding assay cells were prepared according to the standard flow cytometry protocol and stained with 3 µg/mL of coiled-coil AFC for 30 min at room temperature in the dark. Unstained cells were used as a control. For evaluation of ErbB2/Her2 expression A549 and SKOV-3 tumor tissue samples were treated with 50 U/mL of collagenase I and 100 µg/mL of hyaluronidase for 2 hours and filtered through a cell strainer to isolate the cells. After this A549, SKOV-3 cell lines and tumor tissue isolated cells were prepared according to the standard flow cytometry protocol. The cells were stained with Alexa Fluor 488 anti-HER2 mAb (BioLegend, cat.# 324410) for 30 min at room temperature in the dark. Cells stained with Alexa Fluor 488 IgG1k isotype mAb (BioLegend, cat.# 400129) were used as a control.

### SI REFERENCES


1. Gradišar H, Jerala R. De novo design of orthogonal peptide pairs forming parallel coiled-coil heterodimers. J Pept Sci. 2011;17(2):100–6.
2. Choy N, Raussens V, Narayanaswami V. Inter-molecular coiled-coil formation in human apolipoprotein E C-terminal domain. J Mol Biol. 2003;334(3):527–39.
3. Wakankar A, Chen Y, Gokarn Y, Jacobson FS. Analytical methods for physicochemical characterization of antibody drug conjugates. MAbs. 2011;3(2):161–72.
